# Neuronal cultures show bidirectional axonal conduction with antidromic action potentials depolarizing the soma

**DOI:** 10.1101/2021.03.07.434278

**Authors:** JC Mateus, CDF Lopes, M Aroso, AR Costa, A Gerós, J Meneses, P Faria, E Neto, M Lamghari, MM Sousa, P Aguiar

## Abstract

Recent technological advances are revealing the complex physiology of the axon and challenging long-standing assumptions. Namely, while most action potential (AP) initiation occurs at the axon initial segment in central nervous system neurons, initiation in distal parts of the axon has been shown to occur in both physiological and pathological conditions. However, such ectopic action potential (EAP) activity has not been reported yet in studies using neuronal cultures and its functional role, if exists, is still not clear. Here, we show the spontaneous occurrence of EAPs and effective antidromic conduction in hippocampal neuronal cultures. We also observe a significant fraction of bidirectional axonal conduction in dorsal root ganglia neuronal cultures. We investigate and characterize this antidromic propagation via a combination of microfluidics, microelectrode arrays, advanced data analysis and *in silico* studies. We show that EAPs and antidromic conduction can occur spontaneously, and after distal axotomy or physiological changes in the axon biochemical environment. Conduction velocity is asymmetrical, with antidromic conduction being slower than orthodromic. Importantly, EAPs may carry information and can have a functional impact on the neuron, as they consistently depolarize the soma. Thus, plasticity or gene transduction mechanisms triggered by soma depolarization can also be affected by these antidromic APs. Altogether these findings have important implications for the study of neuronal function *in vitro*, reshaping our understanding on how information flows in neuronal cultures.

## INTRODUCTION

A neuron is a highly specialized cell, typically compartmentalized into dendrites, soma, and the axon. The axon is often seen as a mere transmission cable for action potential (AP) propagation but this a very limiting view, partially arising from the technical challenges in recording from the thin axonal branches of vertebrate neurons (Alcami & El Hady, 2019; Debanne, Campanac, Bialowas, Carlier, & Alcaraz, 2011). Recent breakthroughs, made possible by *in vitro* technological developments such as super-resolution microscopy (Chéreau, Saraceno, Angibaud, Cattaert, & Nägerl, 2017), voltage imaging (Peterka, Takahashi, & Yuste, 2011), fluorescence-guided subcellular patch-clamp (Sasaki, Matsuki, & Ikegaya, 2011), microfluidic tools (Holloway et al., 2021; Estrela Neto et al., 2016), or microelectrode arrays (MEAs) (Emmenegger, Obien, Franke, & Hierlemann, 2019) have opened new insights into axonal signal conduction and generated a renewed interest in axon physiology. Accordingly, accumulating evidence shows that the computational repertoire of the axon is much more complex than traditionally thought (for reviews see (Alcami & El Hady, 2019; Bucher & Goaillard, 2011; Debanne et al., 2011; Sasaki, 2013; Traub, Whittington, Maier, Schmitz, & Nagy, 2020)).

After the seminal work by Hodgkin and Huxley on axonal propagation (Hodgkin & Huxley, 1952), the AP initiation and its propagation have been extensively investigated (Bucher & Goaillard, 2011). These studies have placed the axon initial segment (AIS) as the main site capable of AP generation in central nervous system (CNS) neurons. Intriguingly, several studies have demonstrated that APs generated at distal sites of the axon, also known as ectopic APs (EAPs), co-occur in diverse types of neurons both *ex vivo* and *in vivo* (Bähner et al., 2011; Bukalo, Campanac, Hoffman, & Fields, 2013; Chorev & Brecht, 2012; Dugladze, Schmitz, Whittington, Vida, & Gloveli, 2012; Pinault, 1995; Sheffield, Best, Mensh, Kath, & Spruston, 2011; Thome et al., 2018). Such studies opened perspectives on neuronal communication beyond the canonical orthodromic signal transmission (Sasaki, 2013; Traub et al., 2020). In vertebrates, the occurrence of EAPs has been associated with pathological conditions where the axon is hyperexcitable (e.g., epilepsy, nerve injury, demyelination) (Gutnick & Prince, 1972; Hamada & Kole, 2015; Pinault, 1995; Stasheff, Hines, & Wilson, 1993), but also with physiological functions, such as synaptic plasticity (Bukalo et al., 2013; Bukalo, Lee, & Fields, 2016), or fast network oscillations (Bähner et al., 2011; Dugladze et al., 2012; Sheffield et al., 2011). Still, as these studies relied on patchclamp paired recordings, or paired field recordings from brain slices, the fine detection and characterization of the antidromic conduction properties (e.g., conduction velocity) could not be attained. Moreover, many uncertainties still hold regarding the precise mechanism(s) of EAPs’ initiation (Alcami & El Hady, 2019; Traub et al., 2020). Hypotheses such as local depolarization mediated by activity in connected (e.g., axo-axonal coupling) (Bähner et al., 2011; Schmitz et al., 2001) or adjacent axons (ephaptic coupling) (Anastassiou, Perin, Markram, & Koch, 2011; Han et al., 2018), as well as stochastic activation of sodium channels in unmyelinated distal thin segments of the axon (Hamada & Kole, 2015; Pinault, 1995) have been proposed, but the characterization and function of EAPs remain elusive. Importantly, it remains to be shown if EAPs/antidromic APs carry usable information to the cell body or if they are simply an electrophysiological glitch. From a functional perspective, a bidirectional flow of activity would certainly increase the complexity, but also the computational power of the axon (Alcami & El Hady, 2019). Learning and plasticity studies in theoretical neuroscience have demonstrated the significant relevance of hypothetical mechanisms that could propagate signals (namely error signals) back to presynaptic neurons (for review see (Whittington & Bogacz, 2019). This is tightly related to the ideas behind the backpropagation algorithm (Rumelhart, Hinton, & Williams, 1986), a central element in state-of-the-art artificial neuronal networks, such as deep neuronal networks.

In the peripheral nervous system (PNS), however, signal initiation and axonal conduction have different characteristics. *In vivo*, the peripheral process of sensory neurons generates APs distally, which propagate antidromically towards the dorsal root ganglia (DRG). The majority of *in vitro* studies have focused, however, on molecular processes, with no electrophysiological characterizations of axon physiology (Black et al., 2019; Nascimento, Mar, & Sousa, 2018). Thus, it remains to be shown if cultured DRGs recapitulate the ability to generate APs at their distal terminals in both physiological and pathological contexts (e.g. after axotomy).

Emerging technologies that allow probing axonal function with high temporal and spatial resolution can help characterize and understand EAPs/antidromic APs. We and others have combined microElectrode arrays and microFluidics (μEFs) to compartmentalize neuronal cultures in well-defined topologies that allow functional readouts (Habibey et al., 2017; Heiney et al., 2019; Lewandowska, Radivojević, Jäckel, Müller, & Hierlemann, 2016; Lopes, Mateus, & Aguiar, 2018), as well as selective manipulations of the different neuronal compartments (Moutaux, Charlot, Genoux, Saudou, & Cazorla, 2018; Estrela Neto et al., 2020, 2016). In particular, μEFs allow the isolation of axons within microchannels, which are aligned over a set of microelectrodes. Thus, μEFs allow for the detection of propagating axonal signals with very high fidelity and temporal resolution in long-term experiments, which are not possible with any other technique (Lopes et al., 2018; Moutaux et al., 2018).

Here, using μEFs and detailed temporal analysis, we report the consistent occurrence of bidirectional axonal conduction in two different *in vitro* models: dissociated hippocampal neurons and DRG organotypic cultures. Via a combination of extracellular electrophysiological recording/stimulation and fast calcium imaging, we show that evoked antidromic events effectively depolarize the soma, anticipating functional roles of antidromic activity. Addressing this frequent occurrence of antidromic APs in both hippocampal and DRG cultures, we explore possible functional roles of these signals under two different conditions: in the pathological context of axonal lesions, we show that EAPs occur after distal axotomy; additionally, we show that physiological changes in the biochemical environment of the distal axon can promote antidromic activity. Finally, we report differences in the velocity of signal propagation, with antidromic conduction being slower than orthodromic.

## METHODS

### Microfluidic design and fabrication

The polydimethylsiloxane (PDMS) microfluidic chambers used in this study followed a design reported on our previous publications (Heiney et al., 2019; Lopes et al., 2018), with minor optimizations regarding the number of microchannels, as well as their length and width. The number of microchannels was reduced to match exactly the number of microelectrode columns (16 microchannels in total). The microchannel width was reduced to 10 μm to isolate fewer axons per microchannel, thus decreasing the complexity of the recorded signals. Moreover, the microchannel length was set to 700 μm to selectively probe axonal function, as microchannels must be 450 μm or greater in length to restrict the axonal compartment neurite access to axons only (Estrela Neto et al., 2016; Park, Vahidi, Taylor, Rhee, & Jeon, 2006). Concisely, the microfluidic chambers were composed of two separate compartments (somal and axonal) interconnected by 16 microchannels with 700 μm length × 10 μm height × 10 μm width dimensions and interspaced by 100 μm. In specific imaging experiments, chambers with longer microchannels (1000 μm length) and 200 μm spacing were also used.

From the microfabricated SU-8 mold, the PDMS microfluidic chambers were produced by replica molding. PDMS was prepared using a 10:1 mix of silicone elastomer and its curing agent (Sylgard 184, Dow Corning) and degassed using a vacuum desiccator. Polymerization was achieved at 70 °C for 3 h, after which the PDMS microfluidic chambers were unmolded and cut. The medium reservoirs were manually punched with a steel biopsy punch (Ø 6 mm). Microfluidic chambers used for cultures with DRG explants were adapted by adding an extra smaller reservoir (Ø 3 mm), which allowed the seeding of the DRG in a central position, closer to the microchannels (as in (E Neto et al., 2014; Estrela Neto et al., 2020)).

### μEF preparation

μEFs were prepared as previously detailed (Lopes et al., 2018). Briefly, planar MEAs (Multi Channel Systems (MCS), Germany) of 252 titanium nitride recording microelectrodes (30 μm in diameter) and 4 internal reference electrodes (organized in a 16 by 16 square grid) were air plasma-cleaned for 2 min. Then, both the MEAs and the microfluidic chambers were briefly submerged in 70% ethanol, allowed to air-dry inside a laminar flow hood, and sterilized by ultraviolet (UV) light exposure. μEF assembly was guided by a stereomicroscope, to correctly align the microchannels with the microelectrode grid. The μEFs were then sequentially coated with poly-D-lysine (PDL) (20 μg/ml) and laminin (5 μg/ml) to promote cell adhesion. Each microchannel encompassed 7 microelectrodes, as these were interspaced by 100 μm (the first and last microelectrodes were at 50 μm of the microchannel entry and exit point, respectively). Thin MEAs (tMEAs) with 200 μm electrode interspacing were used in calcium imaging experiments, due to their compatibility with high-magnification objectives. When using tMEAs, longer microchannels (1000 μm length) with appropriate spacing were used to encompass 5 microelectrodes.

### Cell culture

Experimental procedures involving animals were carried out following current Portuguese laws on Animal Care (DL 113/2013) and the European Union Directive (2010/63/EU) on the protection of animals used for experimental and other scientific purposes. The experimental protocol (reference 0421/000/000/2017) was approved by the ethics committee of the Portuguese official authority on animal welfare and experimentation (Direção-Geral de Alimentação e Veterinária). All possible efforts were made to minimize the number of animals and their suffering.

Primary embryonic rat hippocampal neurons were isolated from Wistar rat embryos (E18). Tissues were dissected in Hank’s Balanced Salt Solution (HBSS) and enzymatically digested in 0.6% (w/v) trypsin (1:250) in HBSS for 15 min at 37 °C. Then, trypsin was inactivated with culture medium containing 10% fetal bovine serum and washed away. Tissue fragments were mechanically dissociated with a plastic pipette and the cells’ suspension was filtered with a 40 μm strainer (Falcon) to exclude remaining tissue clumps. After cell counting, 150k viable cells suspended in 5 μl were seeded in the somal compartment of the μEF. Cells were cultured in Neurobasal Plus medium supplemented with 0.5 mM glutamine, 2% (v/v) B27 Plus, and 1% (v/v) penicillin/streptomycin (P/S). For stimulated emission depletion (STED) experiments, 75k viable cells were seeded on microfluidic chambers (with 700 or 1000 μm-long microchannels) assembled on 22 × 22 mm glass coverslips (#1.5 thickness, Corning). These microfluidic chambers were peeled-off at 1 DIV and neurons were allowed to grow along the coated patterns (i.e., microchannels patterns).

Primary embryonic mouse DRG explants were isolated from wild-type C57BL/6 mice embryos (E16.5). Lumbar DRG explants were removed and placed in HBSS until use. A single DRG explant was seeded in the somal compartment of each prepared μEF. Cells were cultured in Neurobasal medium supplemented with 0.5 mM glutamine, 2% (v/v) B27, 50 ng/ml of NGF 7S (Calbiochem), and 1% (v/v) (P/S).

For the experiments which involved modulation of the DRG’s distal axon biochemical environment, conditioned medium was obtained from bone resorbing osteoclasts. Bone resorbing osteoclasts were obtained via mouse bone marrow cells flushing. Pre-osteoclasts were obtained after 3 days of stimulation with macrophage colony-stimulating factor (M-CSF 30 ng/mL, PeproTech). Adherent cells were then detached and seeded on top of bone slices (boneslices.com, Denmark) in the presence of 30 ng/mL M-CSF and 100 ng/mL receptor activator of nuclear factor kappa-B ligand (RANKL, PeproTech) (Sousa et al., 2016). After 4 DIV, resorbing osteoclasts’ secretome was collected, centrifuged at 140 g, 4°C, 5 min, and stored at −80°C before use.

All cultures were kept in a humidified incubator at 37 °C supplied with 5% CO_2_.

### Viral transductions

Transductions with adeno-associated viruses (AAVs) were performed to live-image neuronal morphology, axonal outgrowth and calcium imaging. For neuronal morphology, ssAAV-1/2-hCMV-chI-EGFP-WPRE-SV40p(A) (8.3 × 10^12^ vg/ml titer) or scAAV-DJ/2-hSyn1-chI-loxP-mRuby3-loxPSV40p(A) (7.2 × 10^12^ vg/ml titer) were added to the somal compartment (0.3 μl per μEF at DIV 1-5). For performing calcium imaging of neurons extending to the axonal compartment, ssAAV-retro/2-hSyn1-chI-jGCaMP7f-WPRESV40p(A) (5.2 × 10^12^ vg/ml titer) was selectively added to the axonal compartment (0.5 μl per μEF at 9-10 DIV).

All viral vectors were produced by the Viral Vector Facility of the Neuroscience Center Zurich (Zentrum für Neurowissenschaften Zürich, ZNZ, Switzerland).

### Electrophysiological recording

DRG explants at 6-11 DIV and hippocampal neurons at 11-25 DIV were used in the electrophysiology experiments. Recording sessions started after 5 minutes of adjustment to recording conditions. Electrophysiological recordings of spontaneous electrical activity were obtained at a sampling rate of 20 or 50 kHz for 5-10 or 1-2 minutes, respectively, unless otherwise specified. All recordings were obtained using a commercial MEA2100-256 system (Multichannel Systems MCS, Germany). Temperature was maintained at 37 °C by an external temperature controller. For long-term experiments (e.g., axonal activity modulation experiments), and whenever imaging was performed concurrently, recordings were performed with the system mounted on an incubated (37 °C) inverted widefield microscope (Axiovert 200M, Zeiss or Eclipse Ti2-E, Nikon) stage supplied with 5% CO_2_.

### Axonal electrical stimulation and calcium imaging

Calcium imaging experiments were performed with hippocampal neurons at 18-21 DIV (at least one week post-transduction with AAV2-retro-jGCaMP7f). Images were acquired by a sCMOS camera Prime 95B, 22mm (Teledyne Photometrics, UK), mounted on a Nikon Eclipse Ti2-E (Nikon, Japan) inverted microscope with a Nikon Achro ADI 10X/0.25NA, a Nikon Pl Apo 20X/0.75NA, or a Nikon Apo LWD λ S 40X/1.15NA (water-immersion) objective. Image acquisition was performed using Micromanager (Version 1.4) at 200 Hz (5 ms exposure), which allowed for the temporal discrimination of soma depolarizations in response to single electrical pulse stimulations. Electrical stimulations were performed using the MEA2100-256 system’s (MCS, Germany) internal stimulator. Per trial, 5 biphasic voltage pulses (−500/500 to −1000/1000 mV, 100 μs per phase) were delivered to the last microelectrode within a microchannel at 0.5 or 1 Hz. To synchronize electrical stimulation, recording, and fast image acquisition, the whole setup was triggered via a transistor-transistor logic (TTL) signal sent at the start of the MEA recording/stimulation protocol.

The resulting videos were median-filtered, regions of interest (i.e., somas, axons) were delineated manually and ΔF/F_0_ traces were calculated in ImageJ using custom macros. ΔF/F traces were exported for analysis in MATLAB 2018a (The Mathworks, Inc., USA).

### Axonal activity modulation

Distal axotomy was performed as previously described (Park et al., 2006). After a baseline recording, the medium from the axonal compartment was removed and stored for future use. Then, a pipette tip was placed at the entrance of the main channel and vacuum suction was applied. Axons were severed by the resulting air bubble that passed through the main channel. Then, the stored culture medium was returned to the axonal compartment. Axotomy was confirmed by imaging the whole culture pre- and post-axotomy.

For the selective blocking of activity, a medium pre-mixed with 1 μM TTX, a potent fast voltagegated sodium channels blocker, was added to the desired μEF compartment after removing and storing the original medium. A hydrostatic pressure difference was used to impair the flow of TTX between compartments. In the same recording session, we performed recordings before adding TTX (baseline), after adding to the axonal compartment, and after adding to the somal compartment. At the end of the recording session, the exposed compartments were washed-out with three rounds of fresh medium replenishment and then allowed to equilibrate in the stored medium. A final recovery recording was performed on the following day.

Axonal stimulation through changes in the biochemical environment was tested in DRG cultures. For the modulation of the axon terminals’ biochemical environment, osteoclast’s conditioned medium (CM) was selectively applied to the axonal compartment as in (Estrela Neto et al., 2020). First, a baseline recording of DRG cultures at DIV6 was obtained. Afterward, the medium from the axonal compartment was gently aspirated and replaced by 100 μl of osteoclast’s CM. Posttreatment (0h) recordings (20-30 minutes) were started as soon as the baseline stabilized following liquid flow perturbation (less than 1 minute). Two post-treatment recordings (3h and 24h post-treatment) were performed additionally.

In all experiments, cultures were not moved out of the MEA2100-256 system during the experimental protocol of the day.

### Immunolabeling

Hippocampal neurons were fixed at 6-7 DIV for STED imaging. Half the media was carefully aspirated and replaced by 4% PFA (2% final concentration) for 20 min at room temperature (RT). Then, the fixative was washed-out with three rounds of PBS 1× and the fixed cells were permeabilized with 0.1% (v/v) triton X-100 (in PBS) for 5 min and autofluorescence was quenched with 0.2M ammonium chloride (NH_4_Cl, in H_2_O). Non-specific labeling was blocked by incubation with blocking buffer (5% FBS in PBS 1×) for 1 h. Rabbit anti-Tau (GeneTex, cat# GTX130462, 1:1000) primary antibody diluted in blocking buffer was incubated for 1h at RT. After three washes with PBS 1×, incubation with secondary antibody (anti-rabbit Alexa Fluor^®^ 488, Molecular Probes^®^, Thermo Fisher Scientific, 1:500) and 0.3 μM phalloidin 635P (cat# 2-0205-002-5, Abberior GmbH, in PBS) for actin staining, was performed for 1 h at RT. After three washes with PBS 1×, coverslips were mounted in 80% glycerol and sealed.

### Stimulated emission depletion (STED) imaging and analysis

STED imaging was performed in an Abberior Instrument ‘Expert Line’ gated-STED coupled to a Nikon Ti microscope with an oil-immersion 60x 1.4NA Plan-Apo objective (Nikon, Lambda Series) and a pinhole size set at 0.8 Airy units. The system features 40 MHz modulated excitation (405, 488, 560 and 640nm) and depletion (775nm) lasers. The microscope’s detectors are avalanche photodiode detectors (APDs) which were used to gate the detection between 700ps and 8ns. Typically, STED images were obtained near the entrance and exit of the microchannel patterns. After STED imaging, microchannel regions were fully imaged in a widefield microscope (20× objective) for mapping of the culture topology. This allowed for the precise measure of the STED image localization, in relation to the microchannel length. Axons were identified based on the Tau specific staining and the presence of periodic actin rings within the membrane periodic skeleton (MPS). The diameter of axons focused on the maximum wide plan was measured manually. Per axon, at least 5 measures were acquired perpendicularly to the longitudinal axon axis by connecting the brighter outer pixels (most often actin rings).

### Action potential detection and propagation characterization

Raw signals were band-pass filtered (200-3000 Hz) and analyzed offline using custom MATLAB scripts. APs were detected by a threshold method set to 4.5-6× (DRG experiments) or 6× (hippocampal experiments) the standard deviation (STD) of the peak-to-peak electrode noise. An AP time was extracted at this surpassing point and no detection was considered for the next 2 ms (“dead time”). Events propagating along a microchannel were identified based on the extracted AP times. Propagation sequence identification and propagation velocity calculation were performed as previously reported (*The membrane periodic skeleton is an actomyosin network that regulates axonal diameter and conduction*, n.d.) and the scripts are available in GitHub at https://github.com/paulodecastroaguiar/Calculate_APs_velocities_in_MEAs.

For the hippocampal culture experiments, a propagating event had to fulfill the following requirements: event detected over the entire microchannel (7 microelectrodes); time delay between electrode pairs lower than or equal to 1 ms (minimum propagation velocity of 0.1 m/s); each AP time isolated, with no neighboring APs in a 4 ms time window. For the DRG culture experiments, due to the decrease in signal-to-noise ratio (SNR) at the extremities of the microchannels, we only considered the 5 inner electrodes of the microchannels for the forward/backward events ratios. The remaining requirements were kept the same. This stringent detection method profoundly reduced the number of possible propagating events, as it eliminated any ambiguity during bursts and excluded sequences with missing AP times on at least one microelectrode **(Supplementary Fig. S1)**. Due to the very high firing frequency observed during bursts, we consider interpretations from such instances as unclear: during bursts, it is very challenging to track specific APs (identity is lost); given the short inter-spike intervals in a burst, the conduction direction can be easily misperceived as being antidromic if the signal delay between electrodes is in the same order of magnitude as the inter-spike intervals (in an electrophysiology equivalent of the stroboscopic effect).

For the propagation velocity calculations, the extracted AP times were further corrected based on the voltage waveforms. Each AP time, originally identified by the threshold method, was subject to a post-detection time correction (within a limited 1 ms window), allowing the AP time to assume the instant of the maximum absolute voltage of the signal trace. The corrected AP times ensured that the propagation velocity was calculated based on the reference APs’ maximum absolute voltage (instead of the instant that the voltage profile crossed the threshold line). Propagation velocities per event were then calculated by dividing the first-to-last electrode distance (600 μm span) by the delay between AP times of the two electrodes (the first and last electrodes in the detected sequence).

Analysis of possible temporal correlations between AP times in different microchannels was carried out using a method very similar to the calculation of peristimulus time histograms. To assess the time dependence between antidromic APs in a particular microchannel (here called “events”) with the APs in all other microchannels, histograms of the time delays between each event and all APs times were calculated. The AP times were all obtained from a predefined reference electrode (the middle microelectrode) in each microchannel, and each histogram (one per microchannel) was calculated in a limited causal time window of 10 ms. Presumed antidromic APs consistently preceded by an AP in another microchannel would lead, using this method, to a pronounced peak in the histogram centered at the typical time delay between both.

### Simulations of conduction velocity in different axonal morphologies

Analysis of potential causes for asymmetric AP conduction was carried out in NEURON simulation environment (Hines & Carnevale, 1997) using a detailed biophysical model of an axon. The Hodgkin-Huxley formalism was used to describe ion conductances in the axon. Three currents were modeled: fast sodium and rectifying potassium currents responsible for action potentials in hippocampal neurons (Traub & Miles, 1991), and a leakage current supporting the resting potential. Default (original) parameters were used with the exception of: axial resistance Ra = 150 Ω·cm, leakage conductance gL = 0.1 mS/cm^2^, sodium conductance gNa = 0.1 S/cm^2^, and potassium conductance gK = 0.1 S/cm^2^. Simulations were carried out assuming a temperature of 37 °C. Two distinct overall morphologies were considered. In the first, the axon was modeled as a cylindrical structure with a length of 1 mm and a diameter of 0.6 μm, with a spatial grid of 10 μm (100 segments). Different diameter tapering levels were studied by keeping the somal end at 0.6 μm and varying the diameter at the axonal terminal side. Tapering was quantified as the percentage of reduction in diameter in 1 mm distance. In the second morphology, branching was considered. The axon was still modeled as a cylindrical structure but now branching every 250 μm for a full total length of 1 mm on each of the 8 branches. After each branch node the axon diameter was allowed to be reduced. As with tapering, this reduction quantified as the percentage of reduction in diameter in 1 mm distance (%/mm). In both morphologies, conduction velocity was calculated for both propagation directions by providing stimulation (at rheobase level, for 1 ms) at either axonal end.

### Finite-element modeling of the electrical potential inside the microchannels

The tridimensional (3D) finite element model (FEM) geometry replicating the μEF microchannels, was constructed with SOLIDWORKS software (v. 2018, Dassault Systemes SolidWorks Corporation, France). The dimensions’ details for all components of the model are available in **Supplementary Table 1**. Finite element analysis was performed with the AC/DC module of the COMSOL Multiphysics software (v. 5.2a, Stockholm, Sweden). The Electric Current (ec) physics interface was selected, considering a transient time-dependent study. A 3D model physics-controlled mesh was also generated in COMSOL for the constructed MEA 3D geometry model, with the extremely fine mesh option. This model is composed of thirteen different domains, whose description, number and electrical properties are available in **Supplementary Table 2**. Electrical boundary conditions were added to the model: two ground conditions at each of the culture medium domain extremities, and an electric potential boundary condition added to each of the axons’ surfaces. The axon’s boundary condition served the purpose of recreating, in global terms, the biphasic profile observed in extracellular recordings. The derivative of a Gaussian function was used here as the biphasic profile. Instead of stationary, the center of the Gaussian function in the axon’s boundary condition was made to depend on time, introducing a wave motion that propagates the entire axon length.

Two different studies were carried, each corresponding to the AP traveling across one of the two axons. Domain probes were added to all of the electrodes to gather the predicted average electric potential in each time-step for each study solution. The direct solver NUMPS was used for both studies.

### Statistical analyses

Electrodes with a mean firing rate (MFR) of at least 0.1 Hz were considered as “active electrodes”. We defined as “active microchannels” those that had at least one detected propagating event per recording. In the stringent conditions of our conduction detection algorithm, propagating events require consistent readouts in all microelectrodes in the microchannel. Consequently, it is important to note that the number of detected propagating events per microchannel did not necessarily correlate with the firing rate. Moreover, the propagating event detection method excluded most APs within bursts **(Supplementary Fig. S1)**, as these could lead to ambiguous detections of propagation direction. As the total number of detected propagating events could vary greatly across days within the same microchannel/μEF (although their direction flow was generally maintained), we opted for characterizing the ratio of antidromic/orthodromic activity in relative fractions for the analyses of the axotomy and chemical blocking/stimulation experiments. Statistical significance was considered for p < 0.05. All statistical data is presented as mean ± STD, unless otherwise specified. Sample sizes and used tests are indicated in the figures’ legends or in the main text.

## RESULTS

### Tracking signal propagation reveals a bidirectional flow of activity

In this study, we improved on our previous microfluidic chamber design (Heiney et al., 2019; Lopes et al., 2018), by optimizing it for the study of axonal function (details in the Methods section). Importantly, we set the number of microchannels to match the number of microelectrode columns, so that every microchannel would be probed electrophysiologically. The alignment of this microfluidic chamber with 252-electrode MEAs allowed for the probing of 16 microchannels, encompassing 7 microelectrodes each, per experiment **(Fig. 1A)**. This configuration enabled the separation of somal and axonal activity within the same experiment **(Fig. 1B)**. The higher impedance within the microchannels greatly amplifies the otherwise difficult-to-detect axonal signals (Lopes et al., 2018; Pan et al., 2014). Due to the increase in SNR, the mean firing rate (MFR) **(Fig. 1C-E)** and the percentage of active microelectrodes **(Fig. 1F)** were consistently higher within the microchannels.

**Fig. 1.**
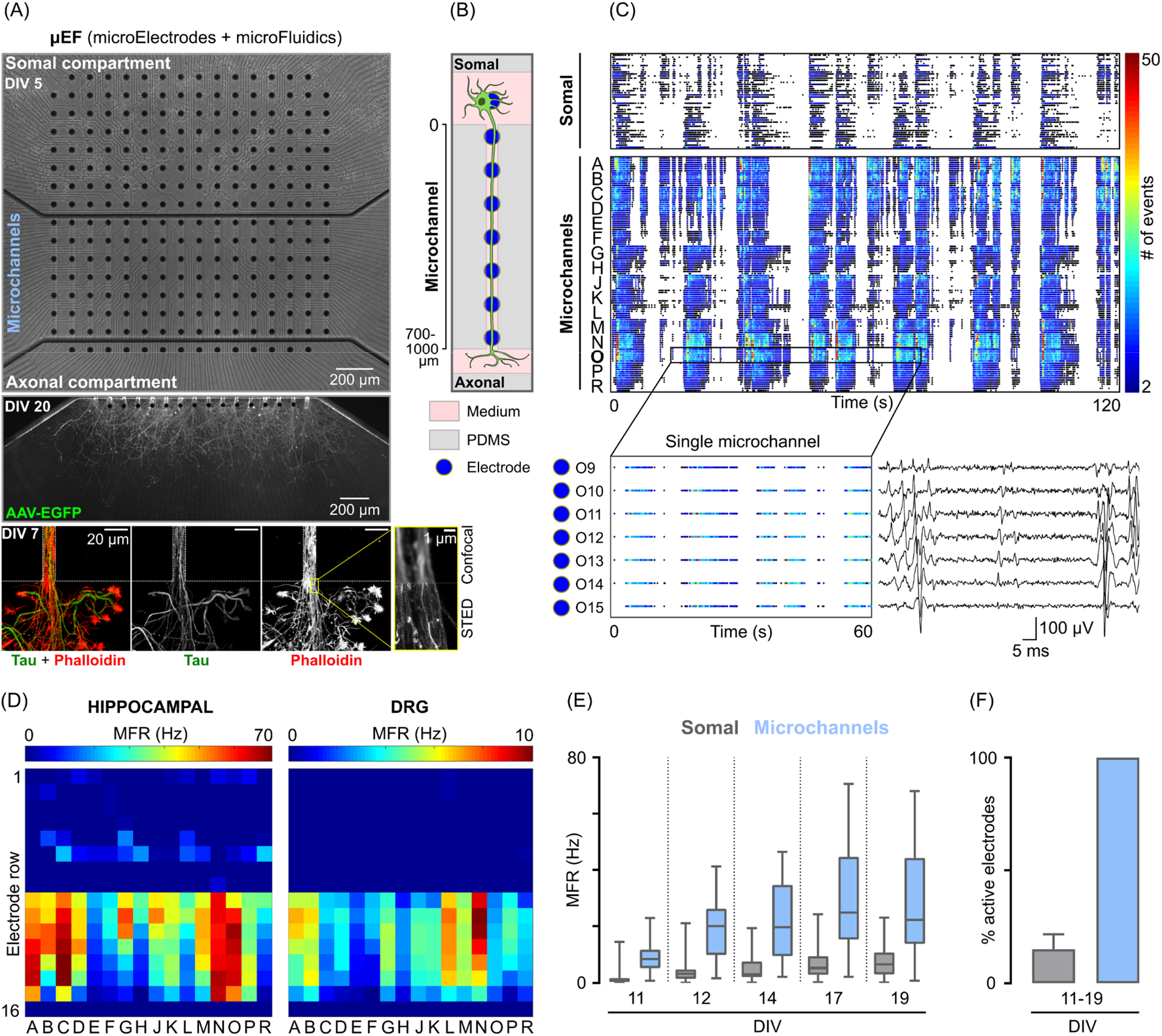
The combination of microElectrode arrays and microFluidics (μEFs) allows tracking of axonal signal propagation *in vitro* with high fidelity. **(A)** Phase-contrast microscopy image mosaic of a hippocampal mono-culture at 5 days in vitro (DIV). The whole microelectrode array (MEA) active area is shown (10× objective, mosaic). A microfluidic device composed of 16 microchannels (10 μm width; 700 μm minimum length) is aligned to include 7 microelectrodes each. Below, an axonal compartment of a hippocampal culture (expressing EGFP) at DIV 20 and the exit of a single microchannel at DIV 7 (stained for Tau and phalloidin) are shown. **(B)** Schematized μEF concept. **(C)** Color-coded raster plots of 2 minutes of activity of all the active microelectrodes of a representative hippocampal culture at DIV19. In total, the activity of 45 microelectrodes from the somal compartment and 112 microelectrodes within the 16 microchannels (A-R) is represented. The inset shows a segment of a single microchannel (O9-15) and an example signal trace. **(D)** Activity maps of a hippocampal and a DRG culture at 19 (same as in (C)) and 7 DIV, respectively. Each pixel corresponds to one recording microelectrode. The mean firing rate (MFR) is color-coded for each microelectrode. Notice that while somal activity can be recorded in the somal compartment of the hippocampal culture, only axonal activity within the microchannels can be recorded in the DRG explant culture. **(E)** Tukey plots of the MFR of all consistently active microelectrodes, during hippocampal maturation, within the somal compartment and microchannels (n = 57 somal microelectrodes, n = 48 microchannels; from 3 independent μEFs). **(F)** Percentage of consistently active microelectrodes, during hippocampal maturation, within the somal compartment and the microchannels (n = 3 independent μEFs).

In the used configuration, single-compartment neuronal cultures (mono-cultures), somata were maintained in a somal compartment and extended their axons along the microchannels to a pure axonal compartment **(Fig. 1A-B)**. We investigated axonal function in two different *in vitro* models: dissociated hippocampal neurons and organotypic cultures of DRG. These different models exhibited marked differences in axonal outgrowth and electrophysiological maturation. Hippocampal neurons’ axons grew through the microchannel within 5 to 7 days **(Fig. 1A)**, while DRG axons took 3 to 5 days. Unlike mature hippocampal neurons, DRG neurons did not fire in bursts but rather exhibited sporadic spontaneous activity. In physiological conditions, this relatively low level of activity also occurs *in vivo* (Black et al., 2019). As DRG explants were placed outside the MEA active area, activity was only recorded in the microchannels **(Fig. 1D)**. This further highlights the importance of microchannels to record axonal activity extracellularly.

We found that in experiments using either dissociated hippocampal neurons or DRG explants’ mono-cultures, a significant number of events propagated from the axonal to the somal compartment - backward propagation **(Fig. 2A)**. This backward propagation completely ceased after selectively adding TTX to the axonal compartment (n = 6 independent μEFs) **(Supplementary Fig. S2)**, supporting the hypothesis that the observed activity initiates in the axonal compartment.

**Fig. 2.**
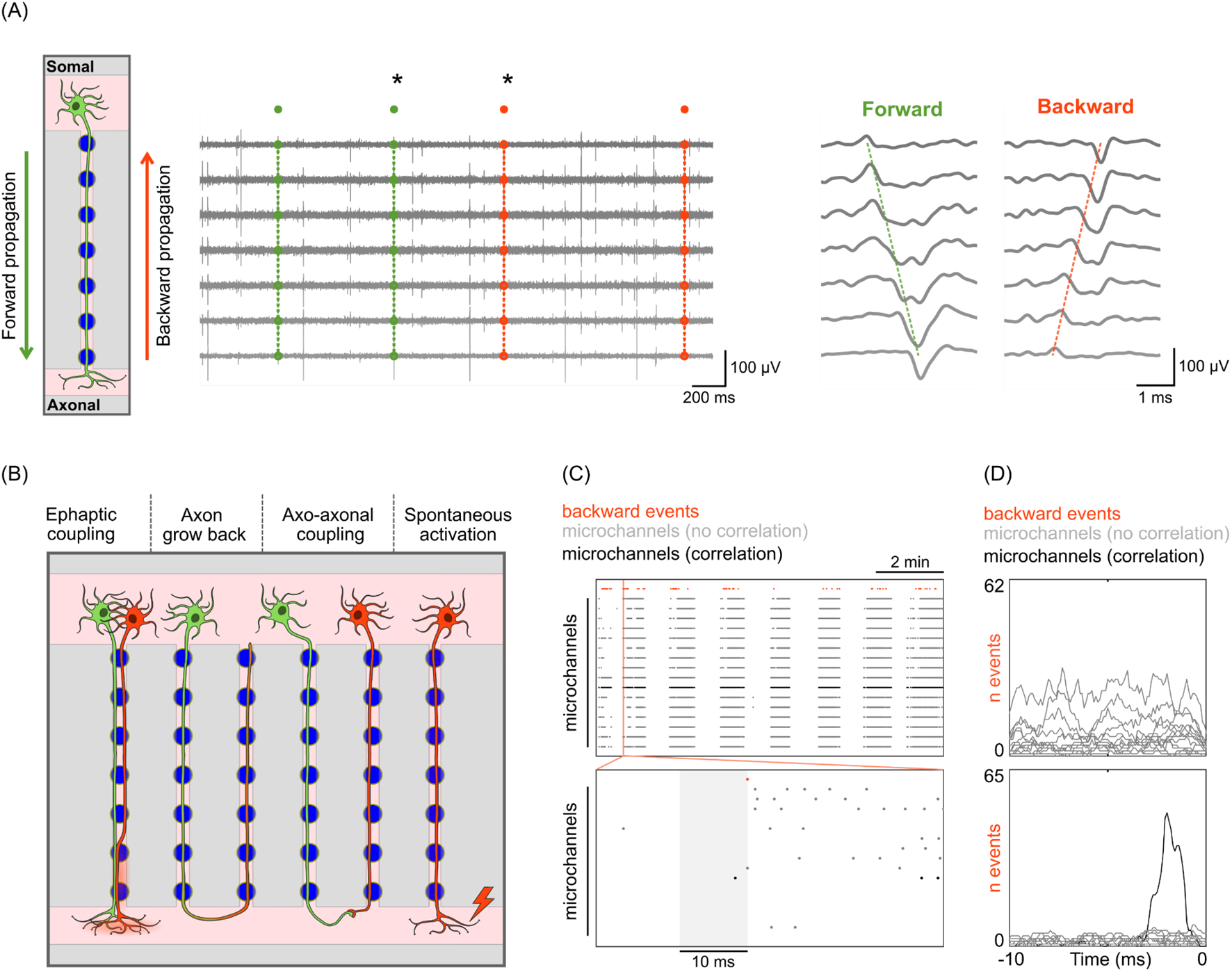
Tracking signal propagation reveals a bidirectional flow of activity which is not explained by axon grow back. **(A)** Schematic representation of the two possible direction flows of signal propagation (forward and backward). Example traces of 3 seconds of activity within a microchannel of a DRG culture at 11 DIV. Detected forward and backward propagating events are marked as green and orange, respectively. The asterisks (*) highlight the two traces expanded in the inset. **(B)** Schematic of the possible causes for backward propagation. Neurons are colored by their propagating direction (forward or backward). Leftmost: nonsynaptic electrical coupling (ephaptic coupling). Center-left: axon grow back to the somal compartment after having extended to the axonal compartment (“U”-turn). Center-right: Axo-axonal coupling via electrical synapse and/or gap junction. Rightmost: Spontaneous activation of the axon distal portion. **(C)** Raster plot of 10 minutes of activity in all microchannels (one electrode per microchannel) from a hippocampal culture at 19 DIV. The first line (in orange) represents all detected backward propagating events (n = 65) from a single microchannel. The zoom-in shows 50 ms of activity in all microchannels. A time window of 10 ms before each backpropagating event was used for isolating possible microchannel correlations (e.g., “U”-turns). **(D)** Pre-backpropagating event time histograms. Examples of no correlation (top) and correlation (bottom; same microchannel as in (C)) are shown.

Different causes can support the observed backward propagation **(Fig. 2B)**. First, the simple case of an axon growing back to the somal compartment after having extended to the axonal compartment (“U”-turn). Additionally, although axons do not find dendritic targets in the axonal compartment, they may establish axo-axonal synapses and/or couple via gap junctions (Schmitz et al., 2001), so that one axon conducts APs orthodromically and the other antidromically along the microchannels. Alternatively, very close proximity between axons in the axonal compartment, combined with high electrical coupling conditions, can originate ephaptic coupling (Anastassiou et al., 2011) where, as with gap junctions or axo-axonic connections, an axon conducts APs antidromically. Finally, this activity may arise from APs initiation caused by activation of sodium channels (stochastic, or not) at the axon distal end followed by antidromic propagation (Hamada & Kole, 2015; Pinault, 1995). The possible origin of the backward propagation was dissected through a set of experiments that selectively manipulated the axonal compartment.

### Backward propagation is not explained by returning axons nor μEF electric artifacts

The μEF design did not prevent axons from growing back to the somal compartment after reaching the axonal compartment (“U”-turns), though these axons would have to elongate for, at least, 1.5 mm. Thus, we tested if the detected backward propagation could simply be caused by axon grow back **(Fig. 2B)**. As the AP activity in returning axons would be temporally correlated with AP activity in another microchannel, we performed time-delay analysis of the backward propagation events against the preceding AP activity in all microchannels **(Fig. 2C)**.

For hippocampal neurons at 11 and 19 DIV (n = 5 independent μEFs), only a minority (<10%) of the microchannels where backward propagating events were detected showed temporal correlations (within 10 ms) with AP activity in other microchannels. An example of neighboring microchannels with a strong temporal correlation is shown in **Fig. 2D**. In DRG experiments at 6 DIV (n = 5 independent μEFs), we found a single pair of temporally correlated microchannels. Moreover, we could not detect a correlation within the same microchannel in any experiment. Thus, the great majority of the detected backward propagation does not emerge from axon grow back to the somal compartment after a “U”-turn. Similarly, the obtained time-delay distributions do not favor the hypotheses of strong axo-axonal or ephaptic coupling as a major cause for the detected backward propagating activity.

A last, but not less relevant alternative hypothesis for the observed backward propagation signals is the presence of electric artifacts in the μEF microchannels. Detailed simulations using FEM, recreating the core geometrical elements of the microchannels **(Fig. 3A)**, were carried out to assess two situations that could be incorrectly interpreted as a backward propagation signal: i) complex electrical behaviors driven by the extracellular currents of forward APs, and ii) influence of axonal activity at the exit of the microchannels on the readout of the microelectrodes. The microchannels’ small cross-section leads to higher impedances and increased recorded signal amplitudes, in accordance with the experimental data **(Fig 3B-C)**. However, the forward propagation of APs inside the microchannels does not generate any unexpected potential readings at the microchannel extremities that could trigger another AP **(Supp. Mat. Movie 1)**. The increased impedance inside the microchannels is also responsible for the fact that axonal electrical activity just outside the microchannel does not generate relevant electrical potential/currents that could trigger an AP in an axon inside the microchannel (i.e., the microchannels do not “channel in” outside currents) **(Fig 3D)**. Finally, the FEM model also excludes the possibility of axonal activity at the exit point of the microchannel being picked up by the microelectrodes inside and incorrectly interpreted as a backward propagation signal. In such case, not only the signal amplitudes in the microelectrodes further away would be extremely low, the signal deflection in all electrodes would be virtually synchronous (as we would be recording the electric disturbance in the medium and not a traveling wave in an axon) **(Fig 3E)**.

**Fig. 3.**
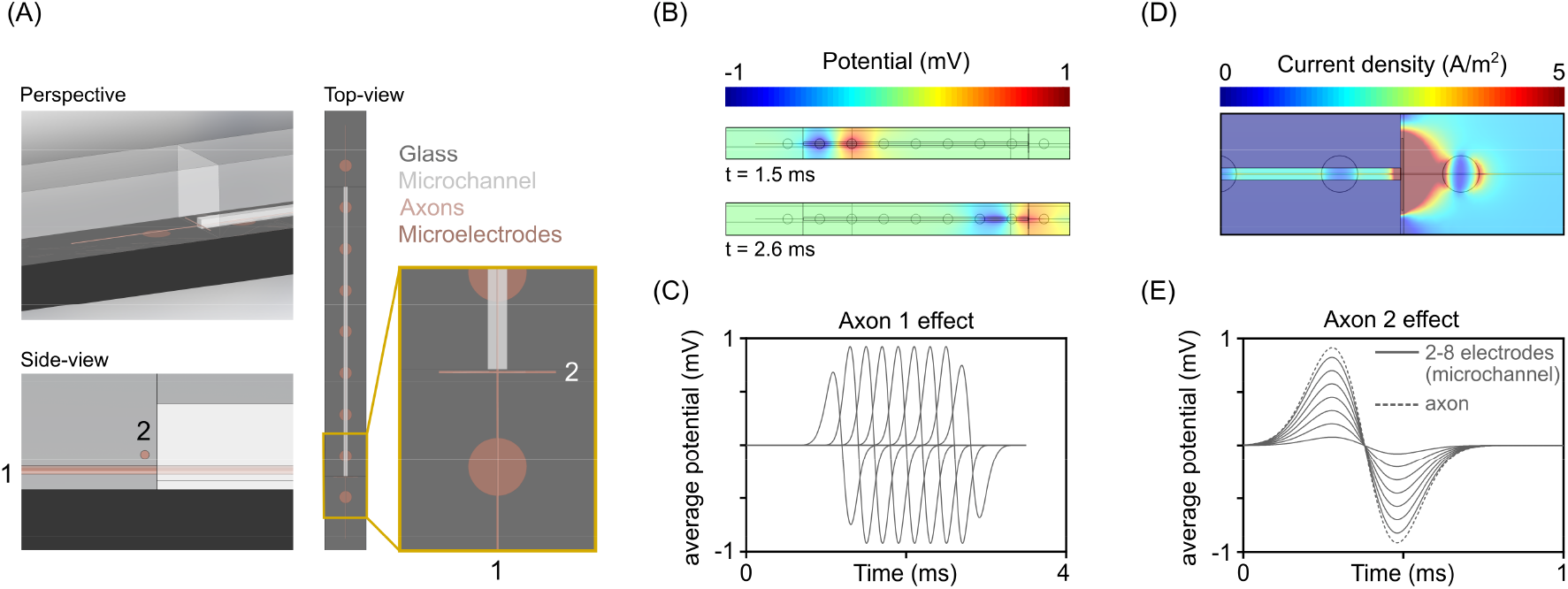
Backward propagation is not explained by electric artifacts. **(A)** Representation of the finite element model (FEM) geometry replicating the μEF microchannels, microelectrodes and axons. Two axons in different positions (axon 1 and 2) were considered: axon inside the microchannel, and axon outside the microchannel and perpendicular to the exit. Three views are presented: perspective, side view and top view with detail on microchannel exit point. **(B)** Simulation of an action potential (AP) propagation along axon position 1. **(C)** Impedance levels inside the microchannel contribute significantly to the ability to record signals from axons. In the model, as well as in experimental data, the amplitudes of the recorded signals are higher in the microelectrodes inside the microchannels. **(D)** Given the higher impedance inside the microchannels, axonal activity at the exit point does not produce current densities that could stimulate axons inside the microchannel. **(E)** The FEM simulations show that (in low noise conditions) electric activity outside the microchannel (axon position 2) can be detected by the microelectrodes inside. However, the signal deflection recorded at each microelectrode would be “simultaneous” (propagation of the electric field in the medium), and would not be confused with an AP.

Concluding, since the vast majority of the detected backward propagating activity does not come from axons growing back, nor electrical artifacts at microchannel entrance, the origin is necessarily the distal parts of the axons. As such, the terminology backward propagation will be replaced by antidromic conduction for the remainder of the manuscript.

### Antidromic action potentials effectively depolarize the soma

For EAPs/antidromic APs to carry usable information to the cell body and have a functional in CNS neurons, they should reach and effectively depolarize the soma. Thus, we tested if eliciting antidromic activity could lead to soma depolarization. For that, we retrogradely transduced hippocampal mono-cultures with ssAAV-retro/2-hSyn1-chI-jGCaMP7f-WPRE-SV40p(A) at 9-10 DIV, which allowed us to perform calcium imaging of neurons extending to the axonal compartment **(Fig. 4A)**. This AAV2-retro variant (Tervo et al., 2016) allows for the robust retrograde expression of the protein of interest. Here, we used it to selectively induce the expression of jGCaMP7f (‘fast’), a genetically encoded calcium indicator (GECI) with fast kinetics and single-AP sensitivity (Dana et al., 2019). As relatively few neurons project axons to the axonal compartment, labeling was sparse even weeks post-transduction **(Fig. 4B)**.

**Fig. 4.**
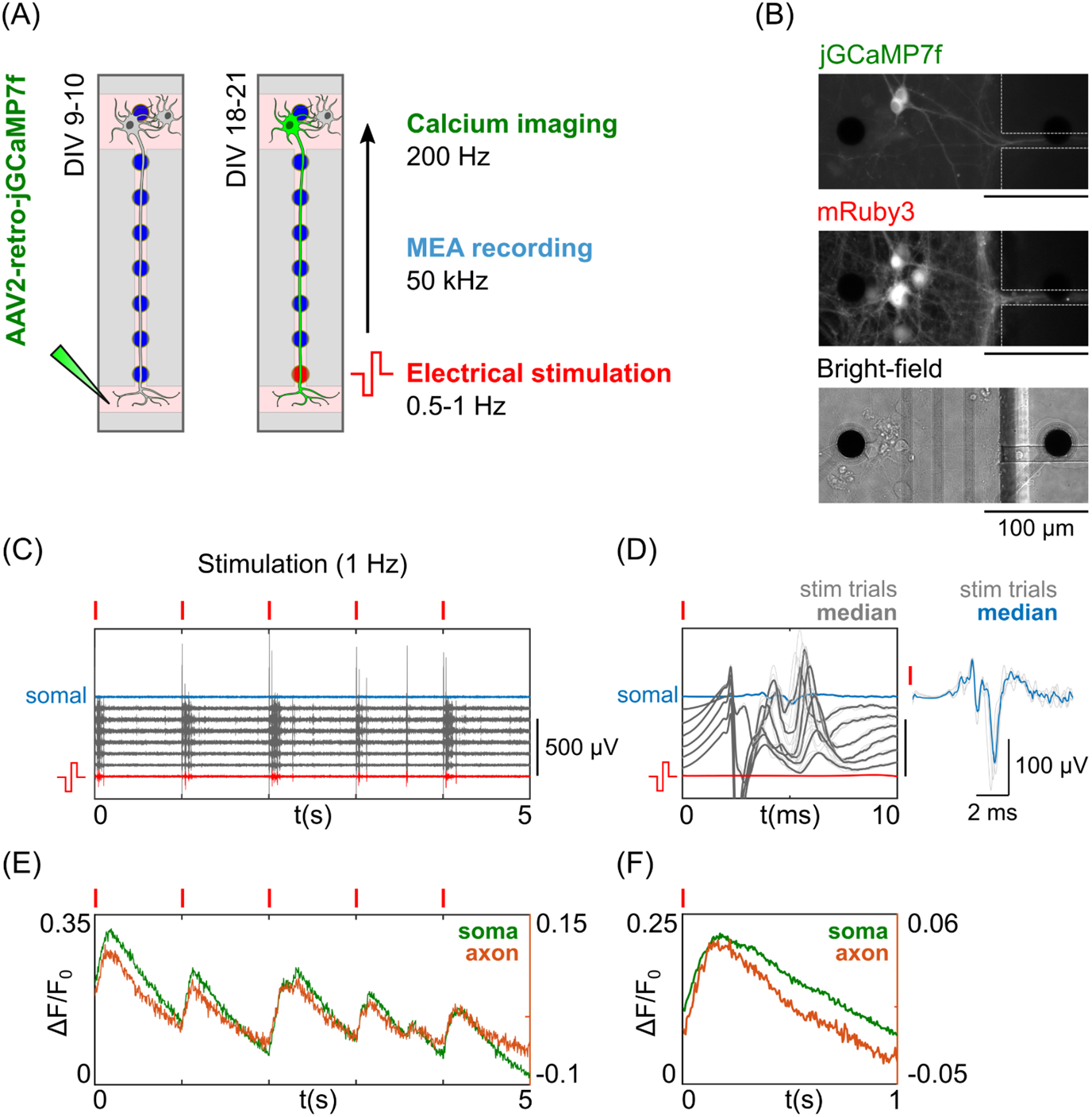
Evoked antidromic activity effectively depolarizes the somatodendritic compartment. **(A)** Schematic of the experimental protocol. Hippocampal neurons extending axons to the axonal compartment were retrogradely transduced at 9-10 DIV with the AAV2-retro-jGCaMP7f. This allowed for distal electrical stimulation of the axon, electrophysiological recording, and calcium imaging of the soma at 18-21 DIV. The last electrode within the microchannel was used for evoking antidromic activity (>600 μm away from the soma). **(B)** Fluorescence (jGCaMP7f and mRuby3) and bright-field imaging of transduced neurons at 21 DIV. **(C)** Electrophysiological traces of 5s of stimulated activity of the microelectrodes of interest, within the microchannel containing the axon (red and gray) and below the soma (blue). For stimulation, 5 biphasic pulses were delivered at 1Hz (−500/500 mV). Red ticks represent the timing of electrical stimulation. **(D)** Overlay of the 10 ms post-stimulation for the 5 trials (peri-stimulus response). **(E)** Calcium imaging traces of the soma and proximal axon’s responses to the distal axon electrical stimulation in (C-D). **(F)** Average calcium response to the 5 stimulation trials.

A standard electrical stimulation protocol was used to elicit antidromic activity on transduced neurons (18-21 DIV), while performing fast calcium imaging and electrophysiological recording **(Fig. 4A)**. For that, 5 electrical pulses were delivered to the last microelectrode within the microchannel of interest (at least 600 μm away from the soma) per trial. The microchannel of interest (i.e., containing the axon of the transduced neuron) could be readily identified via labeling (mRuby3 and/or jGCaMP7f) **(Fig. 4B)**. Antidromic APs were elicited reliably for the range of tested stimulation amplitudes (−0.5/0.5 to −1.0/1.0 V) and frequencies (0.5-1 Hz) **(Fig. 4C** and **Supplementary Fig. S3)**. The resulting stimulation artifact precluded the electrophysiological recording of the immediate responses (within 5 ms post-stimuli) along the microchannel. However, somal depolarizations could be recorded when neurons were near a somal microelectrode **(Fig. 4D)**.

Somatodendritic depolarizations were consistently obtained in response to evoked antidromic APs, as can be seen in the calcium imaging traces **(Fig. 4E-F** and **Fig. S3) (Supplementary Movies 2-3)**. These results were reproduced in several neurons from 6 independent μEFs at 18-21 DIV. With antidromic APs being able to carry information and effectively depolarizing the soma (potentially triggering, for example, protein translation or plasticity mechanisms), the following sections address different physiological contexts capable of triggering/modulating antidromic activity.

### Antidromic action potentials occur after distal axotomy

Spontaneous EAP generation has been shown to occur following axonal injury (Pinault, 1995; Stasheff et al., 1993). Here, we studied the effect of axon lesions on the generation of antidromic signaling. Taking advantage of the high fluidic resistance between compartments, we could subject neurons to distal axotomy >700 μm away from their undisturbed somata. This procedure effectively disrupted axonal projections in the axonal compartment, while sparing the axon proper within the microchannel **(Fig. 5A-B)**, mimicking the conditions of severe axonal lesions (Shrirao et al., 2018). Both before and following axotomy we recorded the spontaneous axonal activity of hippocampal and DRG cultures **(Fig. 5C)**. We will refer to microchannels in which propagating events were detected as “active microchannels”.

**Fig. 5.**
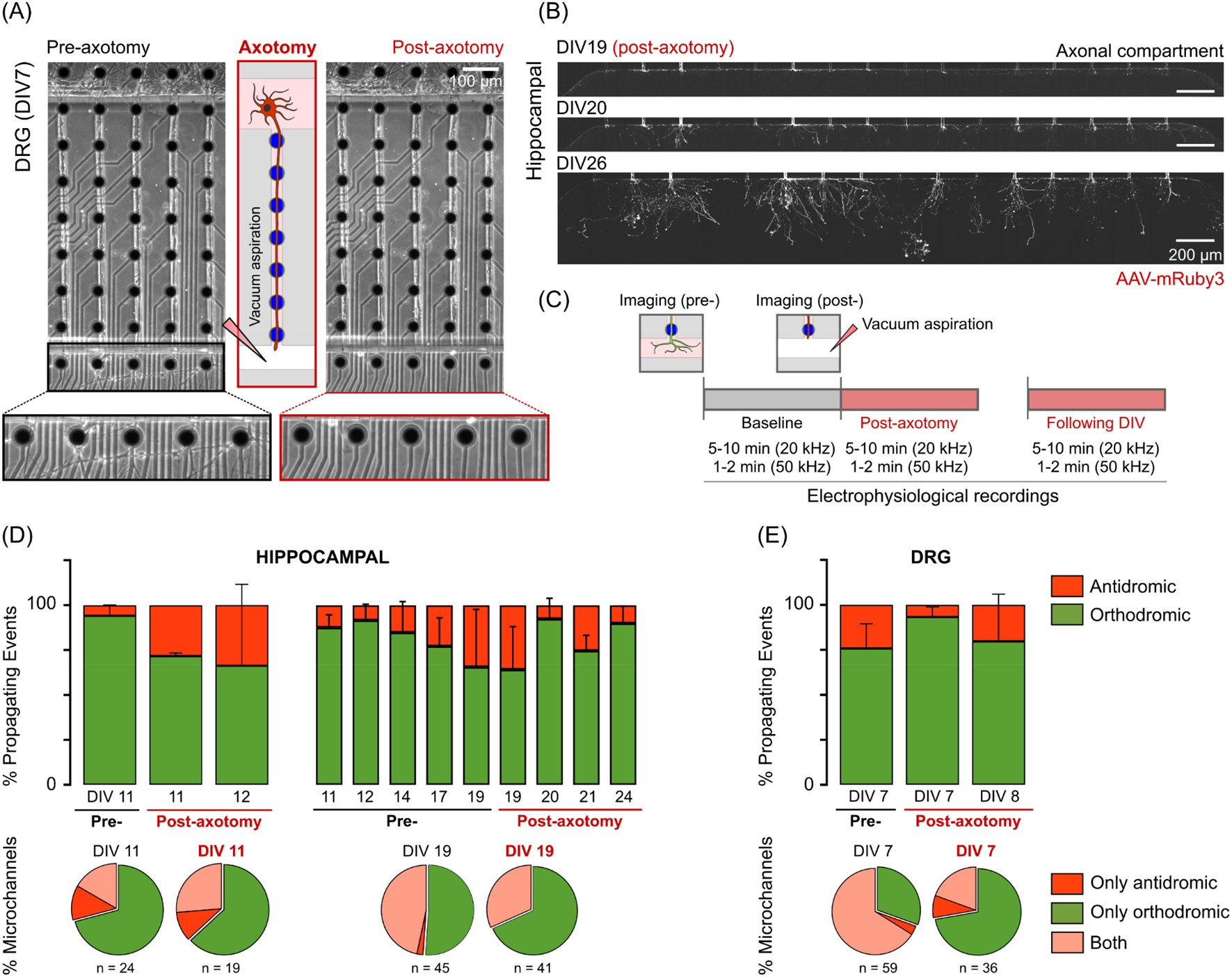
Antidromic activity occurs after distal axotomy in hippocampal and dorsal root ganglia (DRG) cultures. **(A)** Phase-contrast images of a DRG culture at 7 DIV immediately before and after distal axotomy. Axotomy via vacuum aspiration (schematized) effectively disrupted the axonal projections in the axonal compartment, while sparing the axon proper within the microchannel and the somal compartment. **(B)** Mosaic of fluorescence images (expressing mRuby3) of a hippocampal culture at 19 DIV immediately after axotomy, one day after axotomy, and one week after axotomy. **(C)** Diagram of the experimental protocol. (**D)** Bar plots show the percentage of orthodromic and antidromic events in hippocampal cultures (n = 2-3 independent μEF) before and after axotomy (11 or 19 DIV). **(E)** Bar plots show the percentage of orthodromic and antidromic events in DRG cultures (7 DIV) (n = 5 independent μEF) immediately before and after axotomy, as well as in the following day. **(D-E)** Pie charts show the percentage of active microchannels exhibiting only antidromic, only orthodromic, or in both directions’ propagating events immediately before and after axotomy for hippocampal (11 and 19 DIV) and DRG (7 DIV) cultures.

### Hippocampal neurons

Distal axotomy in hippocampal cultures has been explored in different contexts, including in *in vitro* models for post-traumatic epilepsy (Shrirao et al., 2018), an established complication from traumatic brain injury. We studied the electrophysiological effects of distal axotomy in different hippocampal cultures at two stages of maturation: 11 DIV and 19 DIV. Cultures that were subjected to axotomy at 11 DIV (n = 2 independent μEFs), exhibited a significant fraction of antidromic activity immediately after axotomy (28.3 ± 1.7%) **(Fig. 5D)**. We observed, nevertheless, a decrease in the number of active microchannels immediately after axotomy (24 to 19), as well as a decrease in the average number of detected propagating events per minute (from 149 to 61). After axotomy the antidromic events are expected to be triggered at the injury site **(Fig. 5B)**. Immediately after axotomy, 37% of the microchannels (7 out of 19) exhibited antidromic activity. This ratio was very similar at 12 DIV (8 out of 19), although the fraction of antidromic events varied greatly **(Fig. 5E)**. Hippocampal cultures that were subject to axotomy at DIV 19 (n = 3 independent μEFs) showed a tendency to increase the fraction of antidromic activity during maturation. Immediately after axotomy, 32% of the active microchannels (13 out of 41) exhibited a bidirectional flow and a significant fraction of antidromic activity (35.8 ± 24%) **(Fig. 5E)**. The average number of detected propagating events per minute decreased from 104 to 59, immediately after axotomy.

### DRG explants

In sensory neurons *in vivo*, spontaneous AP generation at the site of axonal injury is an important generator of pathological conditions, such as neuropathic pain (Costigan, Scholz, & Woolf, 2009). However, in acute *ex vivo* whole DRG preparations (after lesion), EAPs are rarely encountered, as activity originates primarily in the soma. Typically, ectopic activity needs to be facilitated by applying K^+^ blockers to the recording bath (Amir, Kocsis, & Devor, 2005). Here we studied the effects of distal axotomy on the antidromic initiation of organotypic DRG cultures at 7-8 DIV.

Since most spontaneous axonal activity in DRGs *in vitro* is orthodromic, we analyzed here if ectopic/antidromic activity would emerge after lesion in DRG axon’s distal end **(Fig. 5A)**. At 7 DIV, DRG cultures exhibited a relevant percentage of antidromic events (24.2 ± 13.7%, n = 5 independent μEFs) at baseline (pre-axotomy) **(Fig. 5D)**. We detected antidromic activity in 70% (41 out of 59) of the active microchannels. Immediately after axotomy, we observed a great decrease in the average number of detected propagating events per minute (from 460 to 30). We detected a much smaller fraction of antidromic events after axotomy (6.7 ± 5.7%), which originated from 28% (10 out of 36) of the active microchannels. Together, these results suggest that, while intact, *in vitro* DRG axons generate and conduct antidromic activity; however, immediately after axotomy, although many injured axons are silent, some antidromic activity occurs. Most probably, this activity is the result of AP initiation at the compromised axonal membrane **(Fig. 5A)**. After axotomy, axons may degenerate or, instead, undergo a cytoskeletal reorganization (e.g., axon blebs, neuromas), which affects sodium channel dynamics. This may explain the variability of responses that we observed at DIV 8, when the fraction of antidromic events in the axotomized cultures varied greatly between 0 and 57% **(Fig. 5D)**.

Overall, these results show that EAPs/antidromic activity occur after distal axotomy in both hippocampal and DRG cultures. Interestingly, the two models behaved in opposite ways: DRG explants showed a significant portion of antidromic activity at baseline that decreased after injury, while hippocampal cultures tended to increase the fraction of antidromic activity after injury.

### Modulation of DRG explants’ antidromic activity with osteoclast’s conditioned medium

Previous studies have shown that osteoclastic activity produces changes in the biochemical microenvironment that activate the nociceptors of the innervating DRG nerve terminals (Hiasa et al., 2017). We took advantage of the μEF’s fluidic compartmentalization to selectively stimulate DRG axons with the secretome of osteoclasts cultured in mineralized substrates (active-resorbing osteoclasts) **(Fig. 6A)**. We have previously reported that axonal stimulation with osteoclast’s CM greatly increases the overall levels of axonal activity of DRG explants at 6-7 DIV (Estrela Neto et al., 2020), without assessing conduction direction. Here, we tested if this increase in axonal activity level was accompanied by an increase in the fraction of antidromic activity.

**Fig. 6.**
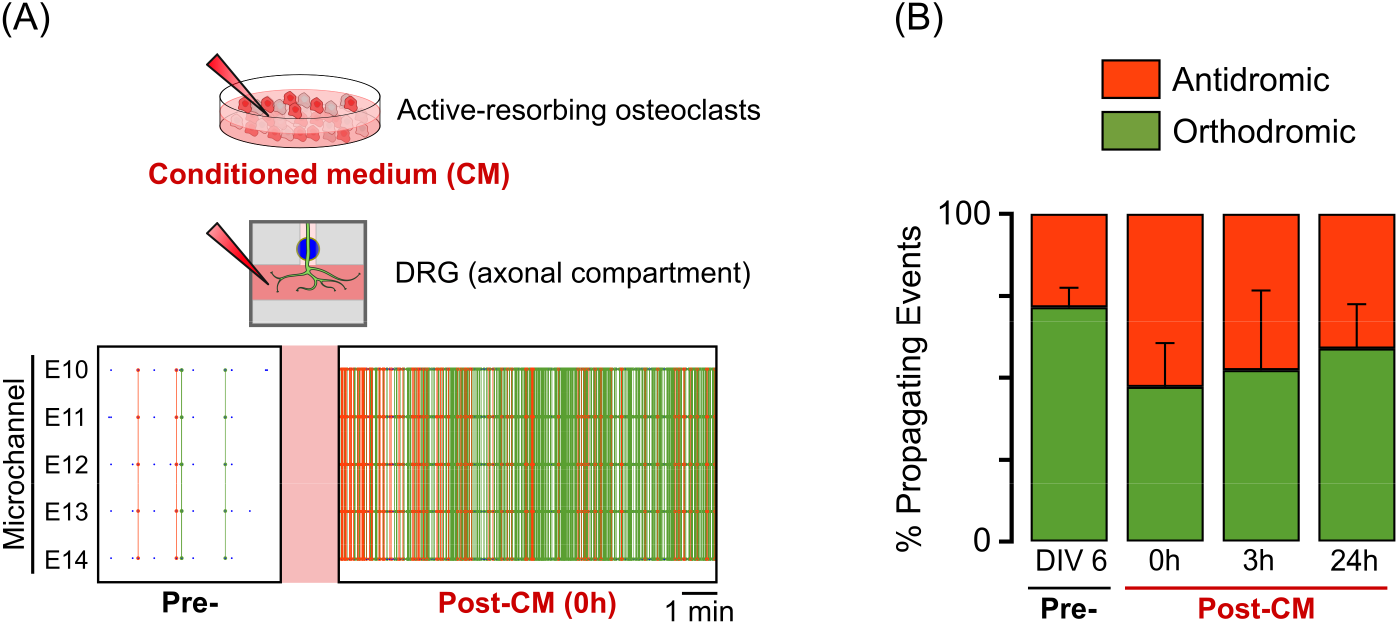
Biochemical stimulation of the dorsal root ganglia (DRG) distal axon modulates antidromic activity. **(A)** Schematic of the experiment of DRG stimulation with osteoclast’s conditioned medium (CM). A representative example of the activity of a microchannel pre- and post-exposure to the active-resorbing osteoclasts’ CM is shown. Orthodromic/Antidromic propagating events are marked as green/orange. **(B)** Bar plot of the percentage of antidromic and orthodromic events in DRG cultures pre- and post-exposure to the CM (n = 3 independent μEFs) at different timepoints (post-0h, post-3h and post-24h).

At 6 DIV, DRG cultures exhibited a relatively low fraction of antidromic events (28.4 ± 5.9%, n = 3 independent μEFs). Immediately after distal axonal stimulation with osteoclast’s CM, this fraction increased to 52.8 ± 13.4%. The following recording time points (3h and 24h post-CM) showed a tendency for this fraction to return to baseline levels, which may be the result of CM’s metabolization. At 7 DIV (24h post-CM), the fraction of antidromic events was 41.1 ± 13.6% **(Fig. 6B)**. These results indicate that distal axon stimulation with osteoclast’s CM can boost the initiation of antidromic activity. Importantly, by modulating antidromic initiation in a more physiologically-relevant model than commonly-used chemical stimulants (e.g., KCl), these results also show that μEFs may be used to study the physiological generation and function of antidromic activity in DRG cultures.

### Conduction velocity is asymmetric

AP conduction along the axon is a tightly regulated and modulated process (Bucher & Goaillard, 2011). Biophysical parameters of the membrane (Waxman, 1980), variations in axon diameter (Chéreau et al., 2017; Goldstein & Rall, 1974), or ion channel densities and kinetics (Hu et al., 2009) greatly influence axonal conduction velocity. Here, the high-temporal resolution (20 μs) of the μEF recordings allowed for the precise calculation and comparison of propagation velocity between orthodromic and antidromic APs (details in the Methods section).

In hippocampal cultures (n = 3 independent μEFs) we observed a moderate increase in propagation velocity along maturation in both orthodromic and antidromic events **(Fig. 7A-B)**. At 11 DIV the average propagation velocity per microchannel was of 0.40 ± 0.06 m/s (orthodromic) and 0.37 ± 0.03 m/s (antidromic), while at 24 DIV (after axotomy at 19 DIV) it had increased to 0.45 ± 0.08 m/s (orthodromic) and 0.43 ± 0.09 m/s (antidromic). The orthodromic propagating velocity values were very consistent with previous results obtained from unmyelinated hippocampal neurons (Bakkum et al., 2013; Habibey et al., 2017; Yuan et al., 2020). However, antidromic events were generally slower than orthodromic events **(Fig. 7A-B)**. This can be observed in the trend of the microchannels’ average propagation velocity **(Fig. 7A)** and in the shift in the velocities distributions **(Fig. 7B)**. Although, the difference was as large as 21.3% at 19 DIV (immediately after axotomy), in physiological conditions the difference was below 10%. Propagation velocity for both orthodromic and antidromic events was not correlated with the microchannel’s MFR (Pearson’s correlation, p = 0.49, p = 0.31, respectively), thus was independent of spontaneous firing frequency within the microchannel.

**Fig. 7.**
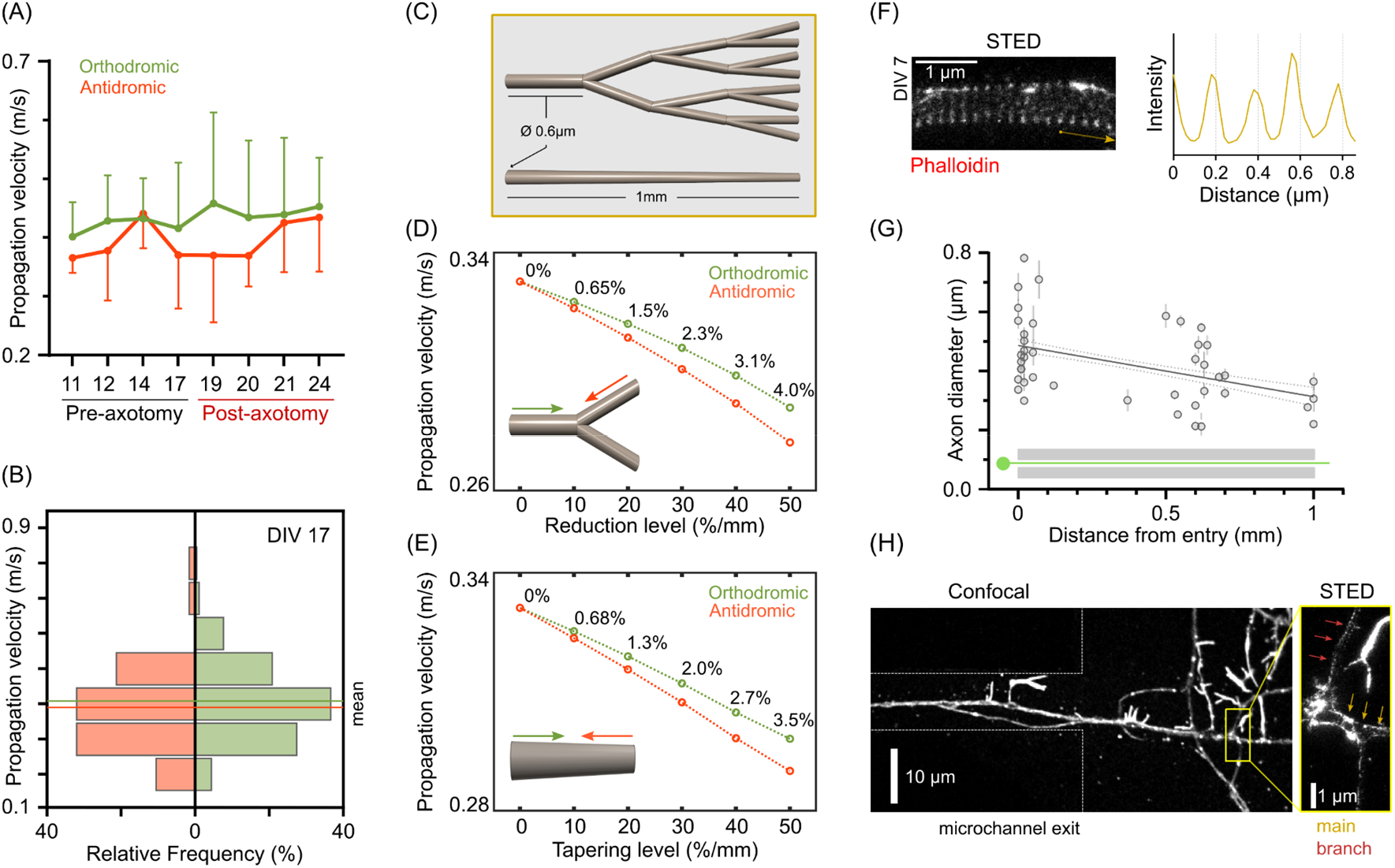
Antidromic conduction is slower than orthodromic conduction due to axon morphology. **(A)** Mean ± STD of the microchannels’ median propagation velocity (orthodromic and antidromic). Distal axotomy was performed at 19 DIV (n = 3 independent μEF). **(B)** Frequency distribution of the propagation velocity for all orthodromic and antidromic events at 17 DIV (0.1 m/s binning). **(C)** Asymmetric axon morphologies in the *in silico* models: step-decrease in axon caliber at branching points and continuous axon tapering. **(D)** Effect of the reduction level (%/mm) in axon caliber at branching points in orthodromic and antidromic propagation velocity. **(E)** Effect of axon tapering (%/mm) in orthodromic and antidromic propagation velocity. **(F)** Representative stimulated emission depletion (STED) microscopy image of an axonal segment (labeled with phalloidin 635) and analysis of actin ring periodicity. The intensity profile was plotted after image processing with a 1-pixel radius median filter. **(G)** Axon diameter as function of distance from the entry of the microchannel (n = 42 axonal segments; from 4 independent experiments) and linear regression. **(H)** Confocal microscopy image of a microchannel exit with a single axon. Inset shows the corresponding STED image of the main axon and its branch.

We used the NEURON simulation environment to test the hypothesis if asymmetry in the axon’s morphology could explain the observed difference. We implemented models to test two different axon morphologies capable of generating asymmetric electrotonic lengths: continuous axon tapering, and step-decreases in axon diameter at branching points **(Fig. 7C)**. We observed that both axon morphologies introduced differences in antidromic and orthodromic conduction velocity, even for mild values of axon diameter reduction after branching or axonal tapering **(Fig. 7D-E)**. The velocity difference was as high as 4%, at 50%/mm diameter reduction.

Beyond the well-described tapering in the soma-to-axon proper transition, little is known on how axon diameter changes along its length (Costa, Pinto-Costa, Sousa, & Sousa, 2018). We performed STED microscopy of axons growing along microchannel patterns to understand if the average hippocampal axon diameter could vary along the microchannel length (up to 1 mm). For the unequivocal identification of axons, we relied on specific stainings (Tau antibody) and the presence of periodic (≈190 nm) actin rings **(Fig. 7F)**, a characteristic cytoskeleton organization (*The membrane periodic skeleton is an actomyosin network that regulates axonal diameter and conduction*, n.d.; Xu, Zhong, & Zhuang, 2013).

Analysis of different axonal segments at 6-7 DIV (n = 42) revealed that axon diameter correlated negatively with distance from the entry of the microchannel pattern (Pearson’s correlation, p = 0.002, r = −0.46, R^2^ = 0.21) **(Fig. 7G)**. The linear regression slope represented a 28-43%/mm (95% confidence interval) decrease in diameter. In some cases, we could measure the diameter of both the main axon and its thinner branch **(Fig. 7H)**. The reduction in diameter at these branching points ranged from 15% to 53% (n = 5 main axon/branch pairs). In summary, STED analysis confirmed the possibility of tapering levels and/or diameter reduction after branching in the order of 40-50%/mm. Thus, the difference in antidromic and orthodromic conduction velocity may originate, at least partially, from these morphological conditions.

## DISCUSSION

The characterization and understanding of EAPs/antidromic activity have been limited by the technical difficulties of performing electrophysiology in the very thin mammalian axons. Recently, high-density complementary metal-oxide-semiconductor (CMOS)-based MEAs have been used to follow AP propagation along the axonal arbors of random neuronal cultures (Bakkum et al., 2013; Yuan et al., 2020). However, due to the very low amplitude of the axonal signals, these studies relied on repetitive electrical stimulation of an axonal segment (usually the AIS) to average out noise (event-triggered averaging) and track signal propagation. In this study, we made use of a powerful combination of methodologies (MEAs and microfluidics) to surpass important technical challenges. This setup allowed us to compartmentalize neuronal cultures and record spontaneous axonal signal conduction events with very high SNR and temporal resolution. Remarkably, μEF recordings of different mono-culture *in vitro* models revealed the unforeseen presence, and significant prevalence, of antidromic APs. These events could not be explained by returning axons (“U”-turns), nor μEF electric artifacts, and were completely abolished after selectively adding TTX to the axonal compartment. These results support the hypothesis of distal initiation of APs in *in vitro* neuronal cultures. Importantly, distal electrical stimulation of hippocampal neurons’ axons consistently led to somatodendritic depolarization, which suggests that antidromic activity may affect neuronal function. We observed similar results with DRG explants that, *in vitro*, show very different axonal conduction characteristics than what is expected from afferent sensory neurons *in vivo*. We explored the possible functional roles of this antidromic activity in the pathological context of axonal lesions and physiological changes of the axonal biochemical environment. We showed that antidromic activity occurs after distal axotomy in dissociated hippocampal neurons as well as in DRG explants. Moreover, the antidromic activity could be biochemically stimulated in physiological conditions by modulating the biochemical environment of the DRG neurons’ axon terminals with osteoclasts’ CM. Finally, we showed that antidromic conduction is consistently slower than orthodromic and that this difference may be explained by axonal morphologies that lead to slight electrotonic length asymmetries.

To date, most studies using μEFs to address axonal function have employed dual-compartment (or co-culture) neuronal cultures of CNS neurons, where signal transmission was assumed to travel orthodromically from the supposed compartment of origin (Gladkov et al., 2017; Moutaux et al., 2018; Pan et al., 2014). Moreover, in the few studies where neurons were seeded in a single compartment, the event detection methods assumed orthodromic propagation from the somal to the axonal compartment (Habibey et al., 2017; Lewandowska et al., 2016). To the best of our knowledge, no study has previously attempted to characterize the signal conduction of DRG cultures. Still, preliminary *in vitro* experiments using DRGs had suggested that most APs propagate orthodromically (Heiney et al., 2019). Our data shows clearly that bidirectional axonal conduction (EAP initiation/antidromic conduction in particular) must be considered when analyzing functional data from both hippocampal and DRG cultures.

Regarding potential functional roles in physiological states, EAP generation at the distal portions of hippocampal neurons’ axons has been reported in studies using *ex vivo* and *in vivo* models, particularly during sharp-wave ripple complexes (Bähner et al., 2011; Bukalo et al., 2013; Chorev & Brecht, 2012; Sheffield et al., 2011). Interestingly, EAP generation is susceptible to conventional synaptic modulation, as somatic excitatory or inhibitory input can facilitate or suppress EAP generation, respectively (Thome et al., 2018). Still, it is not understood in which conditions can EAPs depolarize the soma and have physiological functions. *Ex vivo* studies have shown that backpropagation of EAPs during gamma oscillations can be inhibited by axo-axonic interneurons that target the AIS of CA3 hippocampal neurons (Dugladze et al., 2012). Yet, antidromic activity of CA1 hippocampal axons has been shown to reduce synaptic strength and lead to a widespread downscaling of upstream synaptic weights. Conversely, subsequent synaptic stimulation led to long-lasting synaptic potentiation (Bukalo et al., 2013). More recently, the same authors associated antidromic activity with a rapid downregulation of brain-derived neurotrophic factor (BDNF) mRNA levels (Bukalo et al., 2016), a protein critically involved in canonical forms of synaptic plasticity. Ultimately, these studies suggest that orthodromic and antidromic activity may differentially regulate synaptic plasticity and gene expression. Here, we have shown that EAP generation/antidromic conduction spontaneously occurs in hippocampal cultures. Moreover, we confirmed that evoked antidromic APs consistently lead to somatodendritic depolarization. Given the non-invasive nature of our setup, future studies may assess what is the short- and long-term impact of antidromic APs (spontaneous or elicited) at single-cell and/or network level.

We have also explored the effects of distal axotomy in hippocampal neurons, as in a model for post-traumatic epilepsy (Shrirao et al., 2018), and found that EAP/antidromic activity occurs after axonal lesion. This is in line with previous studies that have shown that, in pathological conditions, EAPs occur during epileptic seizures (Bucher & Goaillard, 2011; Gutnick & Prince, 1972; Stasheff et al., 1993). An important follow-up to our study is the confirmation of the source of the observed antidromic activity in both physiological and pathological conditions. Possibly, in the context of a neuronal network, not all antidromic activity can be traced back to a single cause but results of a combination of mechanisms.

It is not clear if EAPs are as prevalent in CNS neurons *in vivo* as in our *in vitro* conditions, especially in the case of myelinated axons. For instance, myelinated fibers may have evolved mechanistic responses at the nodes of Ranvier that prevent EAP initiation and propagation, as recently suggested (Brohawn et al., 2019). Beyond increasing conduction velocity, myelin may also limit hyper-excitability. In brain slices of mice with cuprizone-induced myelin loss, demyelinated L5 pyramidal neurons were intrinsically more excitable and around 15% exhibited EAP generation. In the study, local application of TTX or K^+^ was sufficient to eliminate or evoke EAPs, respectively, providing compelling evidence for the hypothesis that EAPs may arise from the integration of local environment signals by unmyelinated segments of the axon (Hamada & Kole, 2015). Furthermore, due to the embryonic tissue of origin, our *in vitro* models lack the high prevalence of glial cells observed *in vivo*. Glial cells are known to be key players in the regulation of neuronal excitability, including AP initiation and propagation (Cserép, Pósfai, & Dénes, 2021; Micu, Plemel, Caprariello, Nave, & Stys, 2018; Sasaki et al., 2011). We cannot exclude, therefore, the possibility for an increased prevalence of EAPs when a reduced number of glial cells is present, and the unmyelinated axons extend in a pure axonal compartment.

Most DRG neurons (particularly nociceptive C-fibers) are unmyelinated, however, so far, the study of their axonal physiology has been very limited, as most of our knowledge comes from experiments with myelinated fibers due to their easier accessibility (Black et al., 2019; Nascimento et al., 2018). Here, to the best of our knowledge, DRGs were employed in compartmentalized cultures over MEAs, and selectively manipulated (mechanically and chemically), for the first time. Related studies have obtained functional readouts via calcium imaging or patch-clamp (Huval et al., 2015; Tsantoulas et al., 2013), though these techniques do not have the desirable temporal or spatial resolution, respectively, for assessing axonal conduction dynamics. Tsantoulas and co-workers, in particular, had previously demonstrated somal depolarization in compartmentalized cultures of dissociated DRG neurons after chemical (capsaicin and KCl) or electrical stimulation of the axonal compartment (Tsantoulas et al., 2013). Here, we show that organotypic DRG cultures spontaneously generate APs distally and propagate antidromically both before and after injury, hence recapitulating *in vivo* mechanisms.

Injured sensory neurons exacerbate ectopic activity *in vivo*, a mechanism thought to be an originator of neuropathic pain (Costigan et al., 2009). Here, DRGs’ spontaneous electrical activity and fraction of ectopic activity were diminished immediately after axotomy. It is important to note that silencing following peripheral injury also occurs *in vivo* and it has been proposed as a necessary trigger for axon regeneration (Enes et al., 2010). Likewise, in acute *ex vivo* preparations, EAPs are rarely encountered after lesion, as APs originate primarily in the soma (Amir et al., 2005). Still, the short- and long-term effects of axotomy in DRG axonal function should be the subject of further research, and future *in vitro* studies should explore ways of better mimicking the post-lesion axon environment. Importantly, the antidromic activity could be modulated via distal axon stimulation with osteoclasts’ CM, a physiologically-relevant model of the innervated bone microenvironment (Hiasa et al., 2017; Estrela Neto et al., 2020). The results using DRG explants reinforce the potential of using μEFs as a powerful tool for preclinical testing. Due to their versatility, μEFs could help elucidate conduction dynamics in physiological and pathological conditions.

Further characterization of the bidirectional signal conduction with high-temporal resolution recordings revealed an asymmetry in conduction velocity. This difference could be replicated in computational models of two different morphologies that introduce an asymmetry in axon diameter: axon tapering and diameter reduction at branching points (mimicking propagation into/from higher-order small-diameter collaterals). STED imaging analysis confirmed the occurrence of 36%/mm reductions in average axon diameter, or up to 50% at branching points. Due to the inherent difficulties in assuring axonal identity along the full microchannel length (up to 1 mm), we cannot currently ascertain if this decrease in average diameter is due to axon tapering or branching alone. Moreover, it is not clear if the asymmetry in conduction velocity - which is only dependent on the direction of propagation - has functional implications *per se*.

Importantly, axonal branching has been shown to introduce axonal delays with functional implications, since the main axon temporal activity pattern transforms into multiple patterns in its varying -length and -diameter branches (reviewed in (Debanne et al., 2011)). Concomitantly, related *in vitro* studies have found marked differences (as high as 5-fold) in orthodromic conduction velocity along the length of unmyelinated axon arbors, presumedly due to diameter inhomogeneities (Bakkum et al., 2013; Habibey et al., 2017; Yuan et al., 2020). Here, we reconcile these studies with experimental evidence for axon diameter reduction along the length of microchannel patterns.

Taken together, our results reshapes our understanding on how information flows in neuronal cultures by showing that EAPs/antidromic conduction occur in *in vitro* models of hippocampal and DRG neurons. Several studies have been trying to impose unidirectional outgrowth of axons via complex physical/chemical patterning (reviewed in (Aebersold et al., 2016; Holloway et al., 2021)). Our study suggests that unidirectional axonal outgrowth may not necessarily lead to unidirectional information flow, a finding with important implications for the “brain-on-chip” field. It remains to be shown if these reported mechanisms hold for *in vivo* conditions, but with *in vitro* models being used extensively in conditions where network dynamics play an important role (e.g., plasticity, neuronal circuits), acknowledging the prevalence of these antidromic APs is of fundamental importance.

## Supporting information

Supp. Mat. Movie 1

## ACKNOWLEDGMENTS

This work was partially financed by FEDER - Fundo Europeu de Desenvolvimento Regional funds through the COMPETE 2020 - Operacional Programme for Competitiveness and Internationalisation (POCI), Portugal 2020, and by Portuguese funds through FCT - Fundação para a Ciência e a Tecnologia/Ministério da Ciência, Tecnologia e Ensino Superior in the framework of the project PTDC/EMD-EMD/31540/2017 (POCI-01-0145-FEDER-031540). We thank Alla Karpova from the Janelia Virus Service Facility (HHMI Janelia Research Campus) for providing the AAV-retro vector. All the members of the Neuroengineering and Computational Neuroscience (NCN) group for help and critical discussions. NCN alumni, especially Kristine Heiney, for help and critical discussions. Sean Weaver (ETH Zurich) for critical discussions. Hélder Maiato and António Pereira (Chromosome Instability and Dynamics, IBMC/i3S) for support in STED microscopy. We acknowledge the support of the i3S Scientific Platform Bioimaging, member of the national infrastructure PPBI – Portuguese Platform of Bioimaging (PPBI-POCI-01-0145-FEDER-022122). JCM was supported by FCT (PD/BD/135491/2018) in the scope of the BiotechHealth PhD Program (Doctoral Program on Cellular and Molecular Biotechnology Applied to Health Sciences). MA was supported by FCT through the Scientific Employment Stimulus (CEECIND/03415/2017).

## SUPPLEMENTARY MATERIALS

### Supplementary Figures

**Figure S1.**
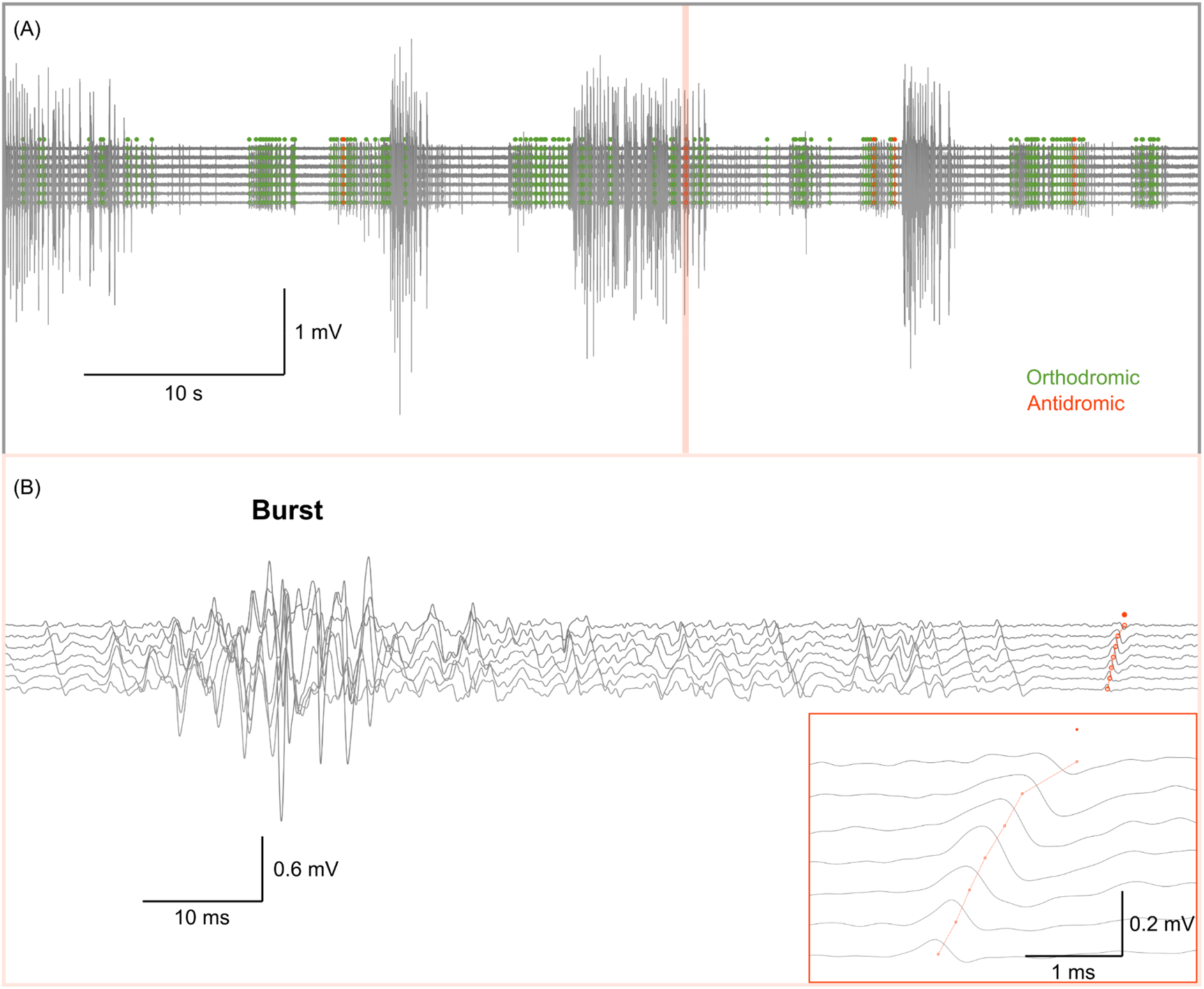
Propagation detection algorithms exclude activity within bursts. **(A)** Traces of 60 seconds of activity within a microchannel of a hippocampal culture at 19 DIV. Note the extracellular activity in the mV range. **(B)** Zoom-in of 100 ms of activity. Due to the stringent detection methods, propagating events are detected outside bursting activity. Inset shows the single detected propagating event (antidromic).

**Figure S2.**
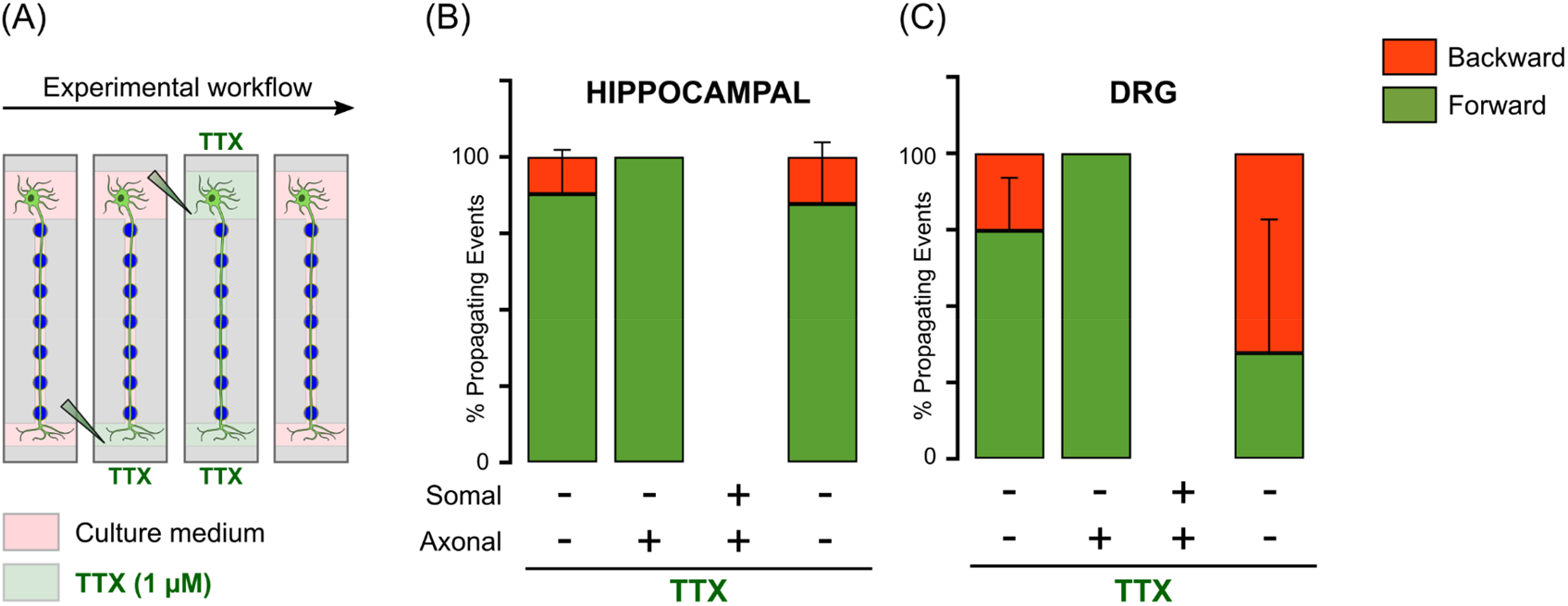
Backward propagating activity completely ceases after selective application of tetrodotoxin (TTX). **(A)** Schematic of the experimental workflow for TTX experiments. After a baseline recording, TTX-containing medium is sequentially added to the axonal and somal compartments. After wash-out, a recovery recording is performed on the next day. **(B-C)** Bar plots of the percentage of backward and forward propagating events in dorsal root ganglia (DRG) (n = 2 independent μEFs at 7 DIV) and hippocampal cultures (n = 4 independent μEFs at 11 or 17 DIV).

**Figure S3.**
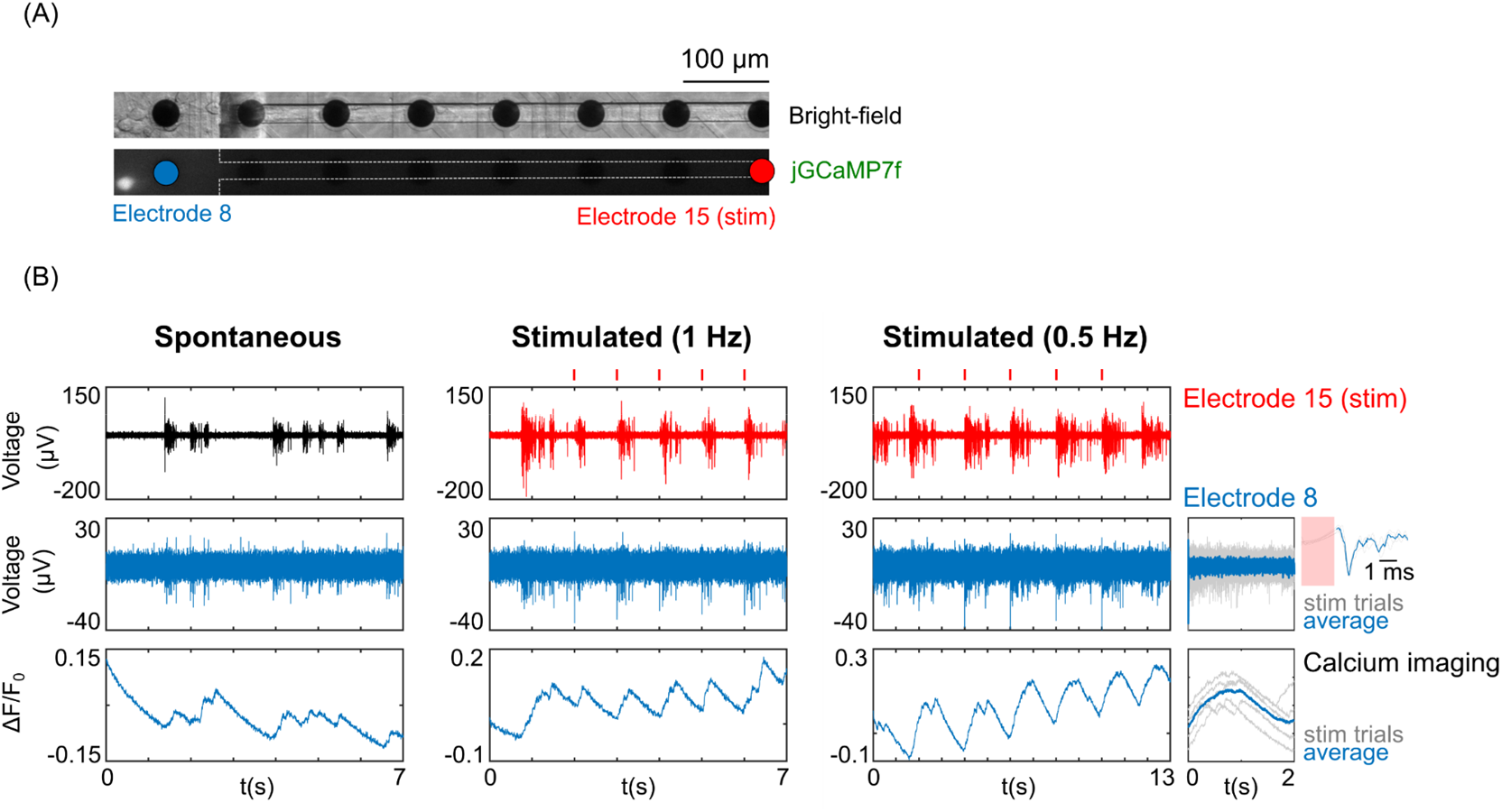
Somatodendritic depolarization is achieved with different stimuli amplitudes and frequencies. **(A)** Bright-field and fluorescence imaging (jGCaMP7f) of a transduced neuron (10× objective). The last electrode within the microchannel (electrode 15) was used for eliciting antidromic activity via electrical stimulation (>700 μm away from the soma). **(B)** Electrophysiological and jGCaMP7f traces of spontaneous and elicited activity. For stimulation, 5 biphasic pulses were delivered at 1Hz (−1000/1000 mV, 100 μs each phase) or 0.5 Hz (−800/800 mV, 100 μs each phase). Red ticks represent the timing of electrical stimulation. Insets show the peri-stimulus response on electrode 8 (closest to soma) and calcium imaging (soma) to 0.5 Hz stimulation.

### Supplementary Tables

**Supplementary Table 1.**
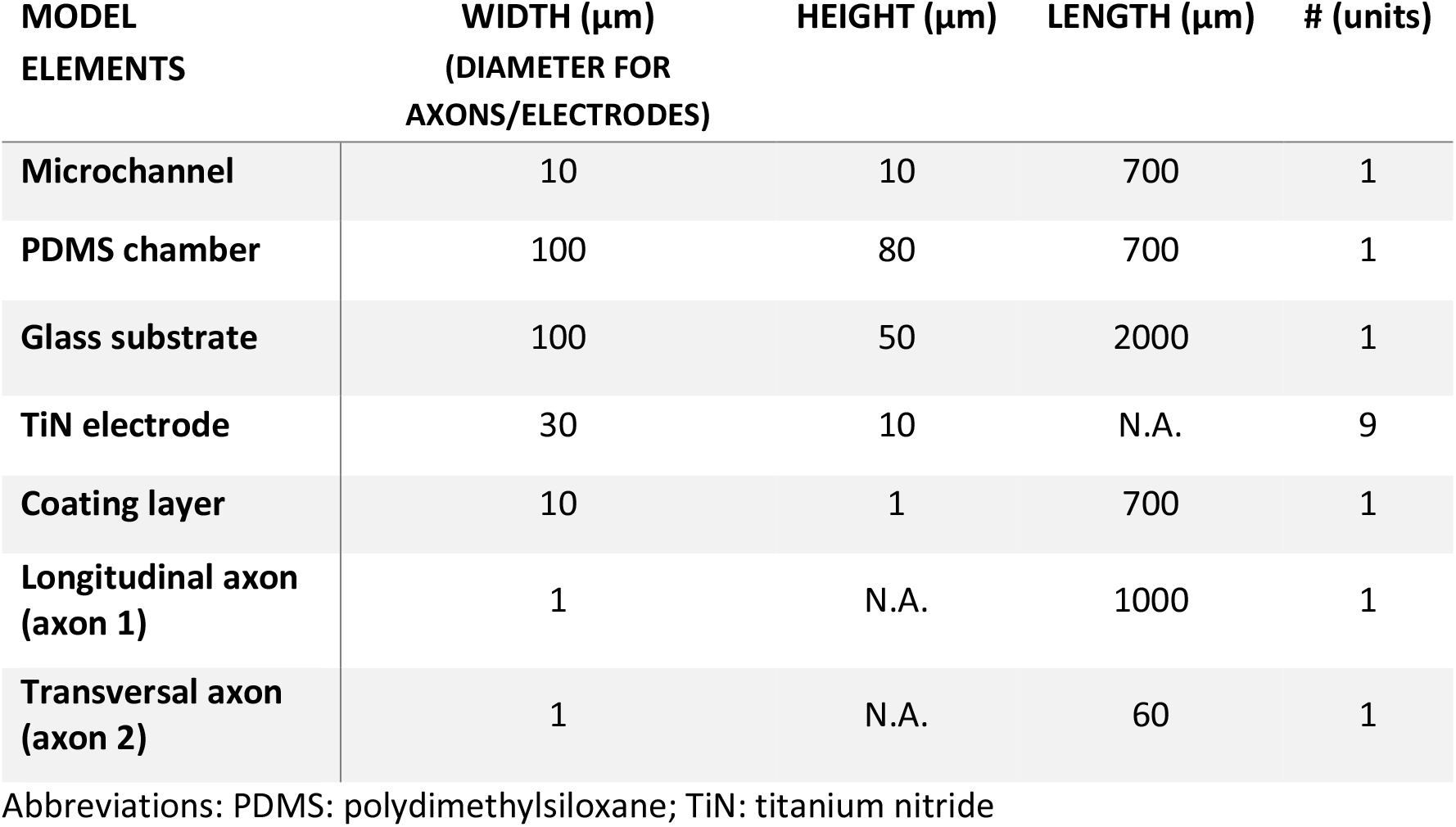
Geometrical details of the 3D finite element model (FEM) replicating the μEF microchannel used in the finite element analysis.

**Supplementary Table 2.**
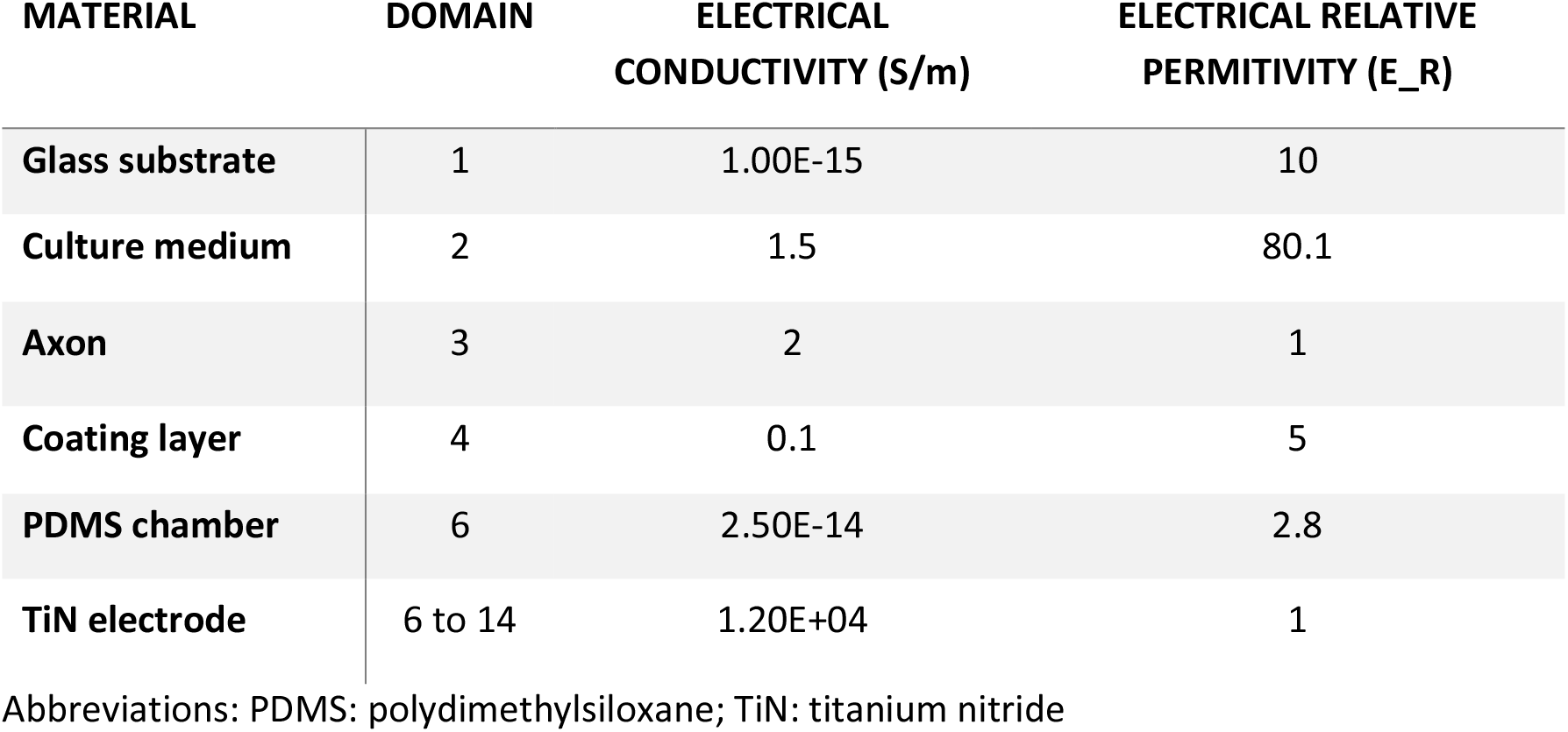
Description of the electrical properties of the different domains of the 3D finite element model.

| MATERIAL | DOMAIN | ELECTRICAL CONDUCTIVITY (S/m) | ELECTRICAL RELATIVE PERMITIVITY (E_R) |
| --- | --- | --- | --- |
| Glass substrate | 1 | 1.00E-15 | 10 |
| Culture medium | 2 | 1.5 | 80.1 |
| Axon | 3 | 2 | 1 |
| Coating layer | 4 | 0.1 | 5 |
| PDMS chamber | 6 | 2.50E-14 | 2.8 |
| TiN electrode | 6 to 14 | 1.20E+04 | 1 |

Abbreviations: PDMS: polydimethylsiloxane; TiN: titanium nitride

### Supplementary Movies

**Supplementary Movie 1** – Finite element model of a propagating action potential inside a microchannel.

**Supplementary Movie 2** – Fast calcium imaging (200Hz; 5s raw movie) of somatodendritic depolarizations in response to distal electrical stimulation (1 Hz; −500/500 mV) (same neuron as in **Fig. 3C-F**). Movie was obtained with the 20× objective.

**Supplementary Movie 3** – Fast calcium imaging (200Hz; 13s raw movie) of soma depolarizations in response to distal electrical stimulation (0.5 Hz; −800/800 mV) (same neuron as in **Fig. S2**). Movie was obtained with the 20× objective.

