## Supplementary figures and images for "Neuronal cultures show bidirectional axonal conduction with antidromic action potentials depolarizing the soma"

### Supp. Mat. Movie 1

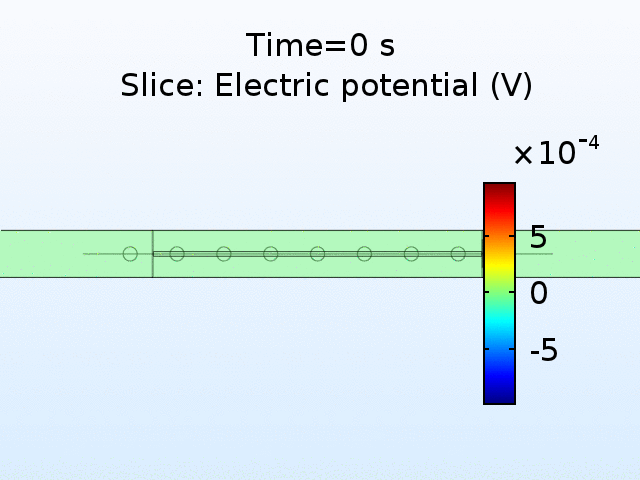
